# Mice dynamically adapt to opponents in competitive multi-player games

**DOI:** 10.1101/2025.02.14.638359

**Authors:** Jeffrey C. Erlich, Mehul Rastogi, Qianbo Yin, Ivana Orsolic, Bruno F. Cruz, André Almeida, Chunyu A. Duan

## Abstract

Competing for resources in dynamic social environments is fundamental for survival and requires continuous monitoring of both ‘self’ and ‘others’ to guide effective choices. Yet most of our understanding of value-based decision-making comes from studying individuals in isolation, leaving open key questions about how animals adapt their strategies during social competition. Here, we developed an ethologically relevant multi-player game in which freely moving mice made value-based decisions in a competitive spatial foraging task. We found that mice integrated real-time spatial information about both themselves and their opponents to flexibly shift their preferences toward safer, low-payout options when appropriate. Analyses of mice behaviour and reinforcement learning agents revealed that these adaptations could not be explained by simple reward learning, but were instead consistent with optimal decision strategies guided by opponent features. Using a dynamical model of decision-making, we found that competition increased choice variability and sensitivity to initial conditions, generating testable predictions for future neural recordings and perturbations. Together, our findings reveal a fundamental mechanism for flexible competitive foraging and propose novel frameworks for quantitatively understanding value-based decision-making in fast-changing social environments.

## Introduction and Results

Our understanding of decision-making is largely built on studies of individual agents making choices in controlled, isolated settings ^1–6^. These investigations have yielded detailed models of how specific factors, such as sensory evidence ^7–9^ or cost-benefit tradeoffs ^10–14^, parametrically shape decisions. Yet real-world decisions are rarely made in isolation; they often depend on the simultaneous and interdependent choices of other agents within the same environment^15^. Classic games like rockpaper-scissors offer a quantitative approach for studying player strategies ^16–23^, but their turn-based structure differs from real-life decision scenarios involving continuous interactions. Meanwhile, existing studies that characterize continuous interactions typically focus on innate behaviours such as prey capture ^24,25^ and courtship ^26–28^. Understanding the biological basis of everyday choices in dynamic and social contexts calls for tractable models of *embodied games*, like tennis or football ^29,30^, where individuals must rapidly and flexibly adjust their value-based decisions in response to the real-time behaviour of others.

To investigate how animals integrate ongoing information about both ‘self’ and ‘other’ to guide choices, we developed the *Octagon*, a novel arena in which freely moving mice harvested reward from eight possible reward patches, either alone or in a competitive social context (Fig. 1a,b, Fig. S1; Methods). On each trial, floor-projected visual stimuli (grating or light triangular slices) indicated the value and location of two active reward patches, each associated with either a high or a low reward (Fig. 1c). The identity and location of the active patches varied across trials, and mice indicated their choice by nose-poking into one of the available options, triggering reward delivery (Fig. 1d). In the social context, only the first response on each trial was rewarded.

**Fig. 1:**
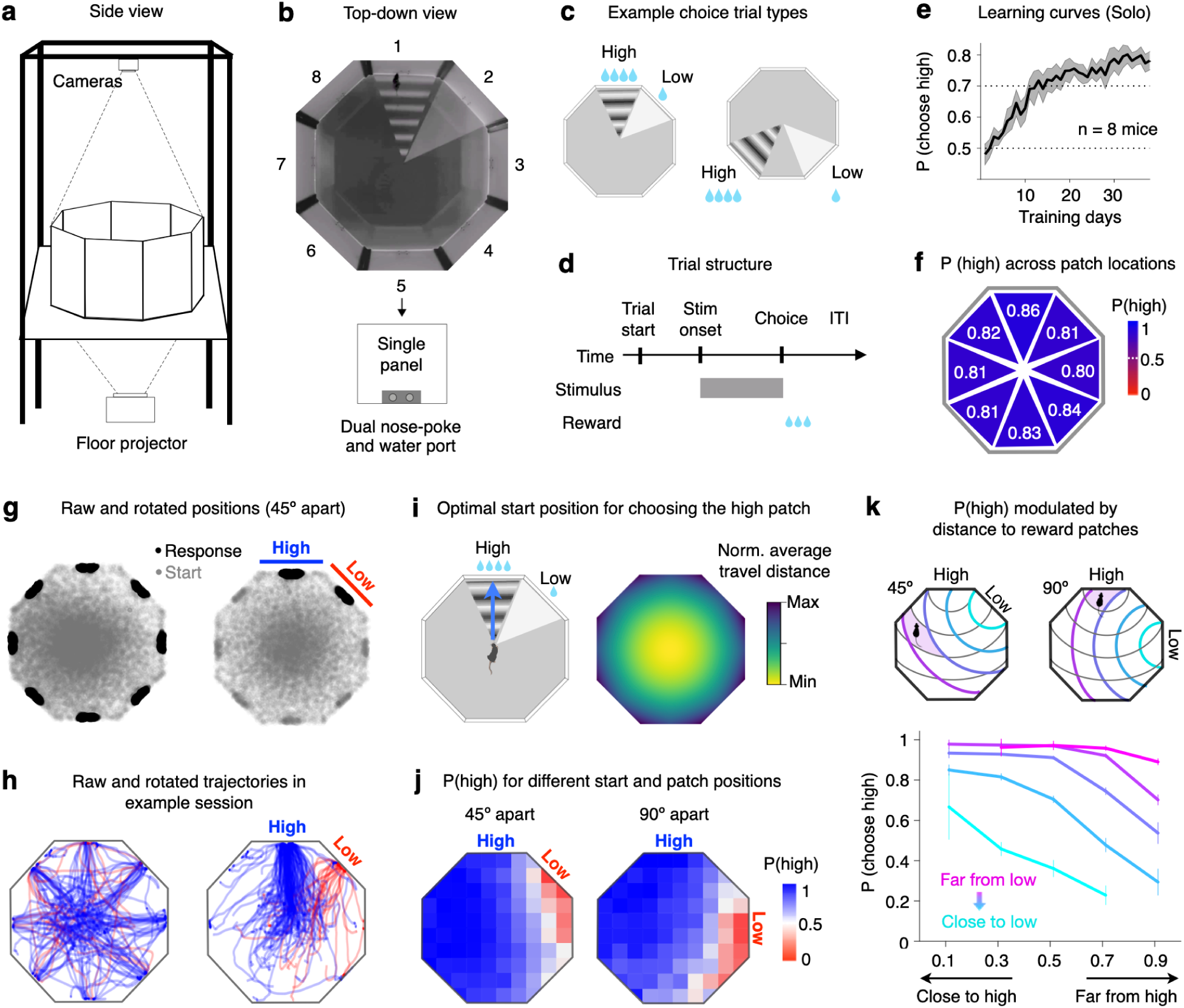
Mice discriminate and generalize patch values in a novel Octagon arena. **a,b**, Side and top-down view of the Octagon arena. Top cameras track animals’ movement; bottom projector displays visual stimuli onto the arena floor. Each of the 8 wall panels has nose pokes and water ports for registering choices and delivering reward. **c**, Example floorprojected visual stimuli that cue the location and value of active reward patches on each trial. **d**, Event timing within a trial. **e**, Solo learning curves of 8 experimental mice, mean *±* s.e.m. **f**, Post-learning preference for the high patch across all 8 wall positions, including trials from 235 sessions and 8 mice. **g**, Left, tracked positions at stimulus onset (gray) and at the time of response (black) for all post-learning solo sessions. Right, rotated positions for visualisation purpose, only for trials where the high and low reward patches were separated by 45 degrees. **h**, Left, mouse movement trajectories from stimulus onset to response for all choice trials (blue, chose high; red, chose low) in an example session. Right, same data but rotated and reflected for visualisation. **i**, Optimal start position to minimize average travel distances (within trials) towards the randomly appearing high reward patch. **j**, Performance across varied start positions in the Octagon, when the high and low patches were separated by 45 degrees (left) or 90 degrees (right). **k**, Top, schematic illustrating how distances to the high and low patches were binned for different trial types. Bottom, the probability of choosing the high patch as a function of varied distances to the high patch (x axis) and to the low patch (coloured lines), error bars are binomial confidence intervals across trials.

The Octagon arena combines the embodied nature of spatial mazes ^31–35^ with the flexibility of virtual reality paradigms ^36,37^, offering an ethological yet quantitative approach for studying strategic interactions between multiple animals. This design provides several key advantages. First, it allows animals to naturally and continuously interact with both the environment and each other during and between decisions. The resulting behavioural variance allows us to parametrically explore how factors such as proximity to reward patches influence foraging decisions. Second, the trial-based structure of the task constrains the timing of patch selection and encourages concurrent decision-making. Third, the ability to flexibly vary the locations of visually cued reward patches across trials dissociates patch value from patch location, helping to disentangle non-spatial foraging decisions from spatial strategies. Finally, the dynamic nature of the task requires animals to actively survey their surroundings on each trial, including the real-time behaviour of their opponent.

### Flexible value discrimination in a dynamic spatial foraging task

Before experiencing social competition in the Octagon, mice were first trained in a solo context to associate distinct visual cues with different reward sizes. Mice learned to discriminate between high- and low-reward patches over a few weeks (n = 8 mice; Fig. 1e); and their post-learning preference for the high reward patch generalised across patch locations (82.2 *±* 0.7%*, n* = 8 locations; Fig. 1f, Supplemental Video 1). The mapping between sensory cues and reward value was counter-balanced across animals, ensuring that the preference was driven by value rather than by specific visual features (Fig. S3a). Response times (RTs) on ‘forced’ trials further revealed value discrimination (Fig. S2). Mice were significantly slower when choosing between two low-reward patches than two high-reward patches (*t*(7) = 4.29*, P* = .0036; paired *t*-test; Fig. S2b). Across sessions, the magnitude of this RT difference was positively correlated with the probability of choosing the high-reward patch on choice trials (*r* = .46*, P* = 1.1 *×* 10^−13^*, n* = 235 sessions; Fig. S2c).

While animals in static environments can adopt simple spatial strategies - such as staying near a high reward location - the dynamic nature of the Octagon encouraged active survey of the arena and higher-order strategies dissociated from specific patch locations. We tracked animals’ continuous positions using Bonsai (Fig. S1c,d; Methods), and quantified how their ongoing behaviours influenced decision preferences as well as their preparatory strategy before each trial. At the time of response, mice were, as expected, located at one of the eight reward patches (black, Fig. 1g left). In contrast, at stimulus onset, their positions were distributed throughout the arena, with a bias towards the centre (gray, Fig. 1g). Animals’ varied start positions allowed us to probe how decision preference was modulated by distance to goals. For visualisation, we rotated and reflected individual trial trajectories such that high and low reward patches were aligned across trials (Fig. 1g,h), and then plotted the probability of choosing the high patch as a function of relative start positions in the Octagon. This revealed a spatial gradient of decision preference: mice generally selected the high reward patch unless they started much closer to the low patch (Fig. 1j). Choice preference for the high patch was negatively correlated with distance to the high patch (*β_dist_*_2_*_high_* = *−*2.53*, P* = 8.65 *×* 10^−108^; GLMM) and positively correlated with distance to the low patch (*β_dist_*_2_*_low_* = 3.41*, P* = 1.76 *×* 10^−269^; Model 2 in Fig. S3c,d, Fig. 1k, Tab. S2). Cross-validated model comparisons confirmed that both distance factors as well as their interaction significantly improved choice predictions (Fig. S3b,c,e, Tab. S5). These findings show that mice developed consistent preference for the high reward option across patch locations; and this preference was modulated by the relative distances to reward patches. As a control, we trained a separate group of mice (‘equal’ mice) to associate the grating and light stimuli with equal rewards (Fig. S4a,b). Unlike the high/low mice, the choices of ‘equal’ mice were dominated by proximity (Fig. S4e-g), further confirming that the preference we observed in the experimental group was driven by value rather than by specific visual features.

Beyond choice preferences, the high/low mice developed a strategy of returning to the arena centre in preparation for each new trial (gray, Fig. 1g). This behaviour emerged without any explicit shaping; and was against rodents’ innate tendency to stay near walls when exploring open spaces ^38^. To test whether returning to the arena centre was an advantageous preparation strategy for dynamic foraging in the Octagon, we simulated the average travel distance from all start positions to randomly positioned reward patches (Methods). For mice trying to select the high-reward patch, starting trials from the arena centre minimised expected travel distance or time on average (Fig. 1i). In contrast, for mice trying to choose the closer of two patches, the optimal strategy involved starting further from the centre, thus increasing the chance of proximity to one of the two options (Fig. S4h,i). Indeed, the ‘equal’ mice did not return to the centre at stimulus onset (Fig. S4c,d). Across all trials, the high/low group started significantly closer to the arena centre (mean normalized distance to arena centre = 0.56) than the ‘equal’ group (mean = 0.78, AUC value = 0.73*, P* = 2 *×* 10^−4^, permutation test). Remarkably, the observed preparatory strategies in the two groups closely matched the optimal solutions for their respective goals, suggesting that mice learned value-sensitive policies that account for distance or time costs in addition to reward.

### Mice integrate self and opponent information to guide competitive choices

After individual mice learned to discriminate patch values in the Octagon, we introduced competitive social sessions interleaved with solo control sessions (Fig. 2a, Supplemental Video 2). In the social context, only the first mouse to report its choice received a reward; the slower mouse received none. Consequently, the effective reward probabilities of the two patches varied dynamically on each trial, modulated by the ongoing behaviours of ‘self’ and ‘other’. How would mice adapt their choices in this competitive context? We considered four possibilities (Fig. 2b): mice might (1) ignore the opponent and behave similarly across solo and social contexts; (2) race to the nearest patch, sacrificing value-based preference, (3) slowly adapt to the changing reward statistics via reinforcement learning without integrating opponent behaviour, or (4) flexibly and rapidly shift strategy by integrating real-time information about both self and other.

**Fig. 2:**
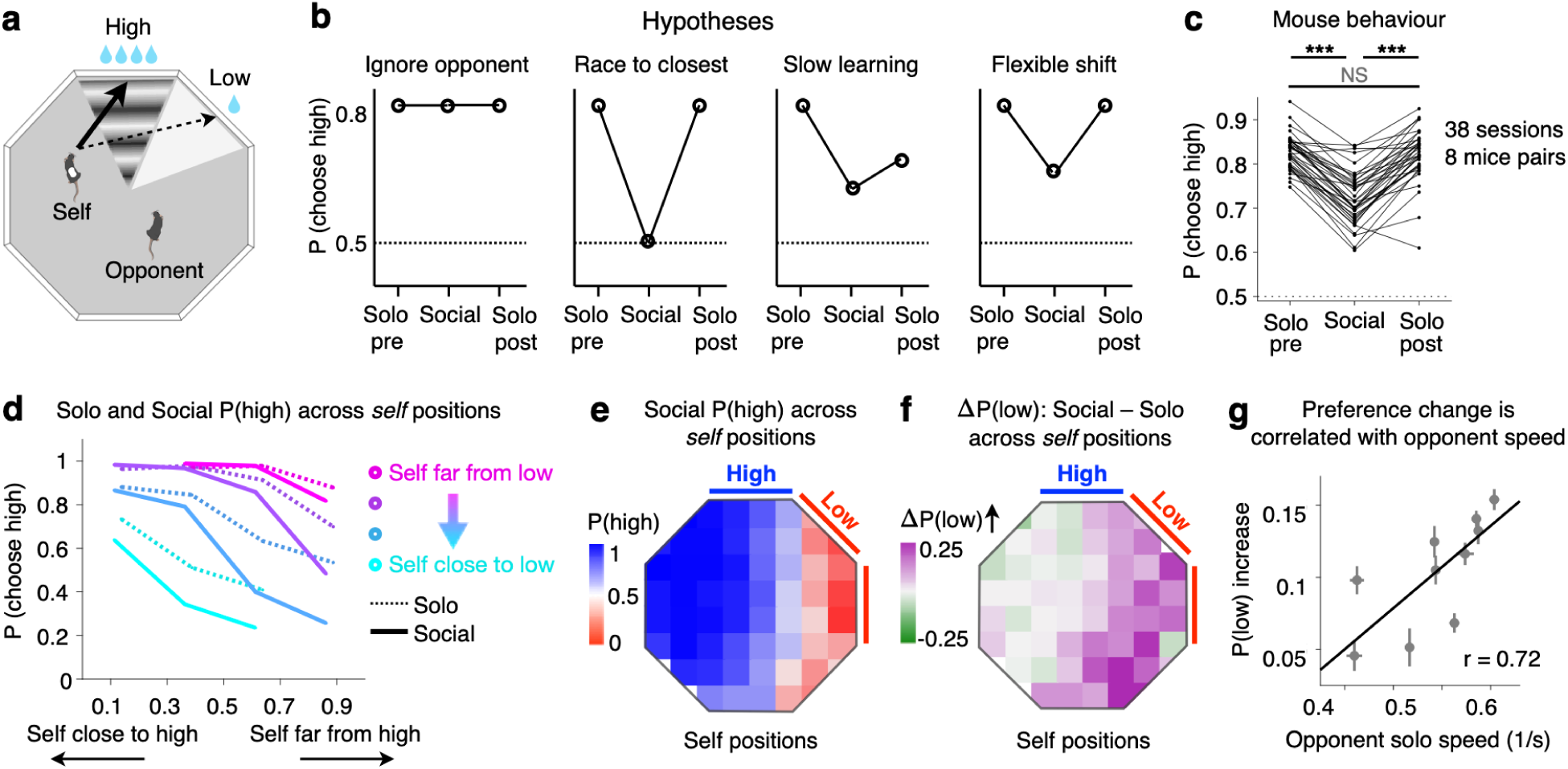
Mice flexibly shift decision preference under social competition. **a**, Schematic for multi-animal competition in the Octagon task. **b**, Hypotheses for how mice would behave across solo and competitive social contexts. **c**, Sessionaveraged change in the probability of choosing the high patch across 38 solo-social-solo experiments from 8 mice pairs. Mice decreased preference for the high reward patch under competition. ***, *P < .*001; NS, *P >* 0.05; paired *t*-tests. **d**, P(high) in solo (dashed) and social (solid) sessions, across varied self distances to the high and low reward patches, similar to Fig. 1k. Self is defined as the winning mouse on each trial. **e**, P(high) across varied self start positions in social sessions (n = 38). **f**, Increase in P(low) from the solo context to the social context across varied self start positions. **g**, Correlation between increase in P(low) in social sessions and speed of the opponent in solo sessions. Mice shifted their strategy more when competing against faster opponents. Error bars are s.e.m. across mice and sessions.

To test these hypotheses, we compared choice preferences across contexts. Mice reliably decreased their preference for the high-reward patch under social competition (10.46*±*0.73% decrease, mean *±* s.e.m., *P* = 7.20 *×* 10^−17^; Fig. 2c), yet still preferred it above chance (*P* = 2.84 *×* 10^−23^), ruling out the first two hypotheses. This preference shift did not change over the course of the session (first half minus second half = *−*0.26 *±* 1.02%*, P* = 0.80; Fig. S5d), nor did it persist into solo sessions on the following day (pre minus post = 0.58 *±* 0.70%*, P* = 0.41; Fig. 2c), arguing against slow, non-social learning. These results suggest that mice flexibly shifted their decision preference across contexts. One possible explanation for this shift is a speed-accuracy trade-off ^39^. In other words, mice might appear less precise (in choosing the high-reward patch) simply because only the faster, winning responses were rewarded in the social context. To test this, we generated RT-matched solo control trials (Fig. S5a,b). Even when RTs were matched across contexts, preference for the high-reward patch remained significantly lower in social sessions (8.58 *±* 0.90% decrease, *P* = 1.18 *×* 10^−10^; Fig. S5c), suggesting that a speed-accuracy trade-off was not sufficient to account for the observed changes under social competition.

What then caused mice to shift away from the high-value option? One possibility is that pursuing the high-reward patch sometimes carried greater risk of losing the competition, particularly when self was far from the high patch and when the opponent was fast. In such cases, choosing the low-reward patch may represent a safer, reward-maximising strategy. Indeed, we observed position-dependent changes in preference (Fig. 2d-f): when self was far from the high patch, mice were up to 24.6% more likely to choose the low-payout option compared to solo sessions. Conversely, when close to the high patch, preferences remained unchanged. Thus, social competition significantly shifted animals’ choices away from the high-payout option in a self-position-dependent manner (*β_social_* = *−*0.62*, P* = 4.45 *×* 10^−27^, GLMM; Fig. S6a-f, Tab. S7). We next asked whether individual opponent features influenced strategy shifts. Across mice pairs, mice shifted more strongly towards the low-reward option when facing faster opponents (*r* = .72*, P* = .02*, n* = 10 opponents; *r* = .32*, P* = .0023*, n* = 88 sessions; Fig. 2g). In contrast, the opponent’s solo value preference did not predict strategy shifts under competition (*r* = .04*, r* = .01*, Ps* = .91 across opponents and sessions). Together, these results suggest that mice adapted their preference flexibly to both their own position and specific behavioural features of their opponent - particularly speed.

Finally, we asked whether mice integrated opponent position to guide their choices on a trialby-trial basis. To avoid bias from analysing only the winner’s behaviour, we included inferred loser’s choices (Methods). For visualisation, we binned self positions and compared P(high) when the opponent was far (Fig. 3a) versus close (Fig. 3b) to the high patch at stimulus onset. Across all trials, mice were more likely to choose the high-reward patch when the opponent was far from the high patch (*β_p_*_2__*_dh_* = 0.27*, P <* 10^−5^, GLMM), even after accounting for self distance effects (Fig. 3a-d; Tab. S11). Moreover, including interaction terms between self and opponent positions improved model fits (Fig. S6g,h; *P* = .003, Tab. S12, S13), suggesting that mice not only tracked opponent position but also adjusted how they weighted their own position accordingly.

**Fig. 3:**
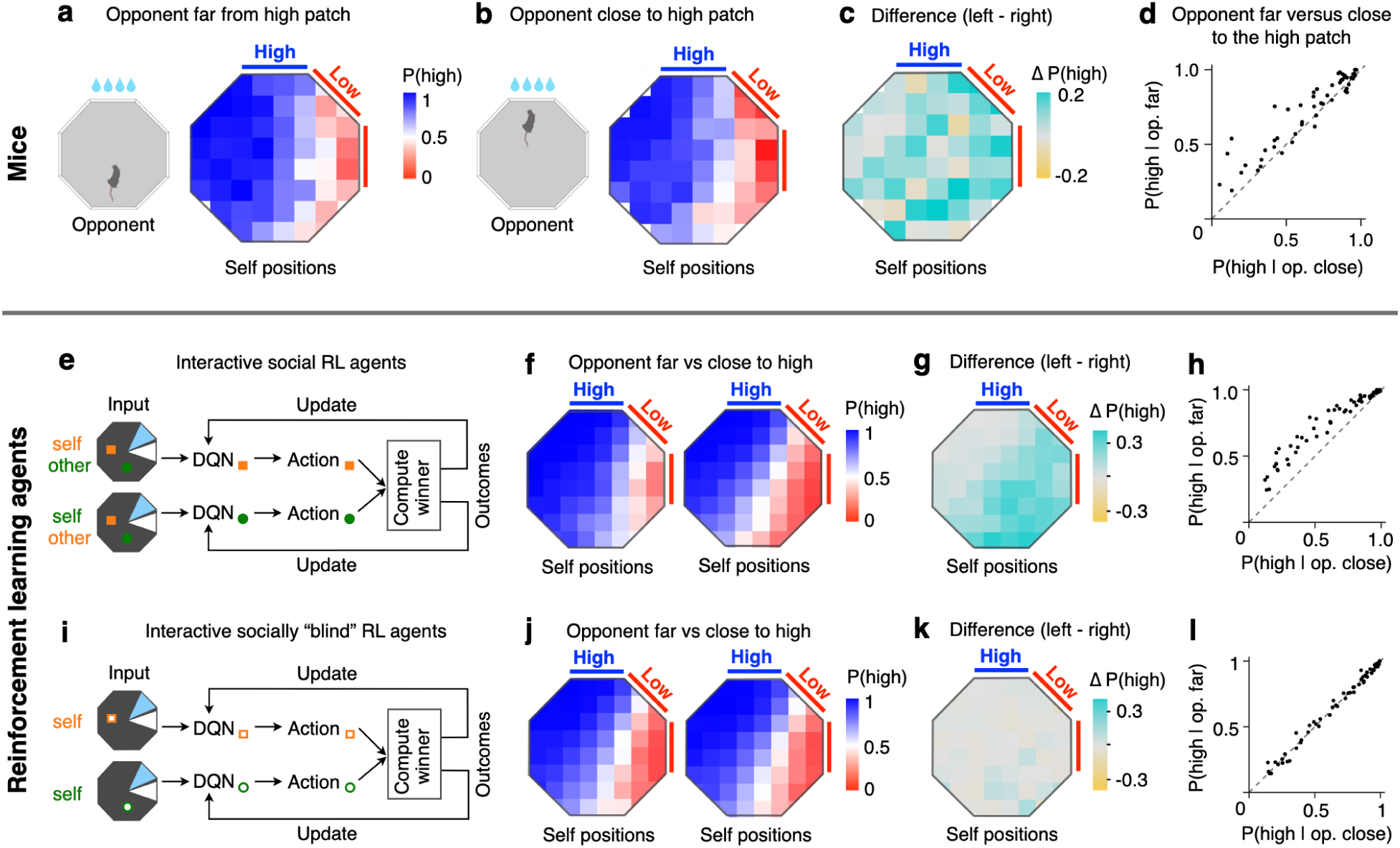
Simulations of multi-agent competition in interacting DQNs. **a,b**, P(high) across varied self start positions in social sessions, plotted separately for when the opponent was far from the high patch (*>* 60% distance, **a**) versus close to the high patch (*<* 40% distance, **b**). **c**, Difference between **a** and **b**. Mice chose the high patch more when the opponent was far from the high patch. **d**, Scatter plot based on data from **a** (y axis) and **b** (x axis), each point represents one specific self position bin. **e,i**, Model architecture and schematic trial structure for two types of DQN agents. ‘Social’ DQN agent (**e**) have access to position information of self and the opponent; whereas ‘socially-blind’ agents (**i**) only have access to self information. Only the winner gets rewarded on each trial (Methods). **f-l**, ‘Social’ agents (**f-h**), but not ‘socially-blind’ agents (**j-l**), show opponent position sensitivity in choice preferences under competition, even with rapid learning (rw = 0.1).

### Interacting artificial agents recapitulate opponent sensitivity in mice

To better understand the strategies used by mice in competitive foraging, we simulated multiplayer games with reinforcement learning (RL) agents trained in the Octagon task. Our goal was to ask whether the flexible preference shifts observed in competing mice could be explained by nonsocial learning, and whether these shifts were consistent with optimal decision strategies guided by opponent features. We implemented two classes of deep Q-network (DQN) agents (Fig. 3e,i, Fig. S7; Methods) ^40^. ‘Socially blind’ agents received environmental and self information, while ‘social’ agents also received opponent information. Both types of agents learned that reward probabilities changed under competition, but only the social agents could use opponent position to guide their decisions. Simulating these distinct artificial agents provided synthetic data to benchmark socially-guided strategies against non-social reward learning.

Consistent with mouse behaviour, DQN agents trained in a solo conditions developed stable value preferences modulated by their distance to reward patches (Fig. S7d; n = 20 agents). After solo training, pairs of ‘social’ or ‘socially blind’ agents were trained in interleaved solo and competitive multi-agent sessions. Agents competed to reach one of the two reward patches, with only the winner receiving a discounted reward and the loser receiving none. We found that simple changes in average preference, such as session-wise decreased preference for the high reward patch under competition, were not unique to social agents. When agents relied on recent experience for learning (high recency weighting, *rw* = 0.1; Fig. S8a; Methods), both social and socially blind agents exhibited rapid, context-specific preference shifts (Fig. S8b,c). Even self-position-specific preference shifts could be mimicked by rapid, non-social learning (Fig. S9).

In contrast, opponent sensitivity, specifically reduced preference for the high-reward patch when the opponent started close to the high patch, was unique to social agents. This pattern held even under rapid learning conditions (Fig. 3f-l). Consistent with mouse behaviours, social agents’ choices were significantly modulated by the opponent distance to the high patch (*β_p_*_2__*_dh_* = 2.1*, P <* 10^−99^; Tab. S15), and this effect varied systematically with self positions in the octagon (interaction *P <* 10^−22^; Fig. 3g, Tab. S16, S17). Socially blind agents showed no such modulation (*β_p_*_2__*_dh_* = *−*0.03*, P* = .59; Fig. 3j-l, Tab. S19, S21), despite showing average preference shifts across contexts (Fig. S8c). Together, these interactive RL simulations confirmed that the opponent-guided strategies observed in competing mice are reward-maximising, and cannot simply be explained by non-social learning alone. They also provide a benchmark and valid null hypothesis for interpreting the flexibility and efficiency of mice’s decision-making in competitive, dynamic environments.

### A dynamical model of flexible decisions under multi-agent competition

Although RL simulations provided a quantitative framework to understand mice strategies, this approach has limitations for probing the neural mechanism underlying flexible decision-making. First, DQNs are black-box models. Dissecting the features and representations that guide their choices would require considerable work which is tangential to our goals here. Second, DQNs lack internal dynamics and stochasticity: their output is a deterministic function of the input, and they do not generate reaction times.

To overcome these limitations and provide a foundation for interpreting future neural recording and perturbation experiments, we turned to well-established dynamical models of decision-making in neuroscience ^41–43^. Inspired by the observation that mice seemed to be ‘drawn’ to the low reward patch when very close to it, we modelled the binary choice between the high and low reward patches as a bistable attractor model ^44^. The model consisted of two mutually inhibitory decision nodes, whose activity evolved based on patch value, self-excitation, noise, and opponent effects (Fig. 4a). To fit the model, we used a combination of global optimisation and gradient descent to find parameter sets that best explained solo and social experimental data respectively. On each trial, the initial state of the dynamical system was based on animal’s distance to the patches at stimulus onset (Fig. 4b). When the animal was closer to the high patch, the high node began near the decision threshold, biasing decisions toward high (Fig. 4e left). When closer to the low patch, the low node began near threshold, leading to fast low-patch decisions (Fig. 4e-right). However, sufficient noise could drive the state to escape the shallower low-patch attractor and settle into the deeper high-patch attractor. Using distance-to-patch as the initial conditions of the dynamical system recapitulated the distance modulation of value preference (compare Fig. 4f with Fig. 1j) without requiring explicit time or action cost as in the DQN model.

**Fig. 4:**
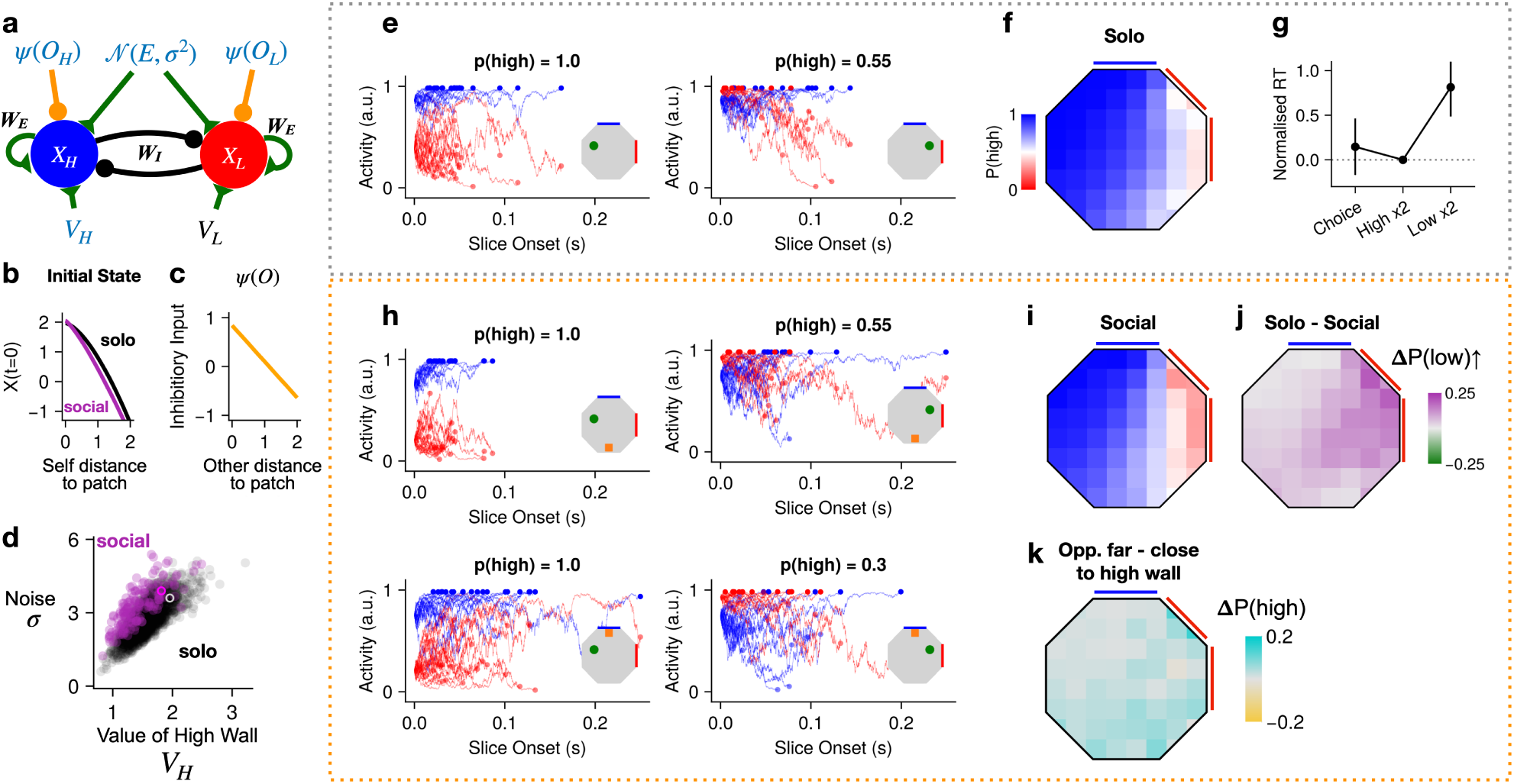
Dynamical model of choice. **a**, Decision dynamics were simulated by a two-node system (*XH*,*XL*) representing the decision to choose the high or low patch. *VH, VL* represent the value inputs to the high and low patch. *WE, WI* represent the weight of self-excitation and mutual inhibition. *ψ*(*O*) represents the influence of the opponent in the social context. *N* (*E, σ*^2^) represent Gaussian noise. Blue parameters were fit to data. Triangles and circles represent excitatory and inhibitory connections respectively. **b**, The initial activity level of the two nodes was determined by the agent’s distance to each patch. **c**, The distance of ‘other’ to each patch modulated the respective node. **d**, Comparison of the noise parameter, *σ*, with the value of the high patch, *VH*, for the top 2.5% of fits to the solo (black) and social (purple) mouse data. White and magenta circles indicate the best solo and social fits, which were used to generate the plots in **e-k**. **e-g**, Simulations generated using the best solo parameters. **e**, Trajectories (*n* = 20 trials) of the solo dynamical system where the agent is close to the high (left) or low reward patch (right). The inset octagon indicates the position of the agent (green circle) and the high (blue) and low (red) patches. The title indicates average P(high) for the 20 simulated trajectories. **f**, Solo preference plot (compare with Fig. 1j). **g**, Simulated reaction times for choice and forced trials (High *×*2 and Low *×*2; compare with Fig. S2b). **h-k**, Simulations generated using the best social parameters. **h**, Trajectories (*n* = 20) of the social dynamical system. The inset shows the trial configuration as in **e** but with the opponent position at slice onset shown (orange square). **i**, Social preference plot (compare with Fig. 2e). **j**, Social - Solo preference (compare with Fig. 2f). **k**, Influence of opponent position on agent choice (compare with Fig. 3c).

Opponent influence was introduced as a distance-weighted inhibitory input to the decision nodes (Fig. 4a,c). When the opponent was far from the patches, the model’s behaviour resembled solo sessions (Fig. 4h-top). When the opponent was close to the high patch, this shifted dynamics and choices toward the low option, especially when self was near low (Fig. 4h-bottom). Compared to the solo model, the best-fitting social model had higher noise and was more sensitive to position and less driven by value (Fig. 4b,d). Together, these mechanisms allowed the model to account for both the self-position-specific preference changes (compare Fig. 4i,j with Fig. 2e,f) and interactions between selfand opponent-position effects (compare Fig. 4k with Fig. 3c,g).

Although the model was fit only to choices between the high and low reward patches, it also captured aspects of reaction time (RT) on other trial types. Simulations of forced-choice trials, where both nodes received equal value inputs, showed that RTs were fastest when choosing between two high-reward patches, slowest between two low patches, and intermediate between high and low (compare Fig. 4g with Fig. S2b). This matched mouse behaviour and suggested that decisions between two low-value options may reflect a process where noise pushed the system back and forth between two shallow basins of attraction.

The model also makes several neural predictions. First, decision-selective neurons should encode abstract value, invariant to the specific patch locations. Second, self distances to patches should be decodable at stimulus onset and decay as the system commits to a decision. Third, opponent distance to patches should also be reflected in decision-related activity. Finally, even when overt choice patterns are identical, the underlying decision dynamics may differ: comparing the left panels of Fig. 4e,h we see that the model predicts 100% choices to the high patch, but the degree of overlap between red and blue traces varies considerably. This suggests that the robustness of the dynamical system, and in turn their susceptibility to perturbation, varies across trial configurations, a testable hypothesis for future causal experiments ^45^.

## Discussion

In this study, we established a novel, freely-moving mouse paradigm to investigate value-based decision-making in a fast-changing, competitive social environment. The dynamic nature of the task encouraged mice to develop flexible and generalizable value-guided strategies across varied trial configurations (Fig. 1). This allowed us to disentangle value from spatial proximity in the solo decision context (Fig. 1j,k), and revealed how animals traded off these factors under competition (Fig. 2c-f). Future work incorporating finer-grained behavioural characteristics, such as head direction, field-of-view, and movement trajectories ^46–48^, may uncover further embodied modulations of decision-making.

Beyond decision preferences, mice developed higher-order strategies such as returning to the centre of the arena before each trial (Fig. 1i), likely to minimize delay between stimulus onset and reward. This is not consistent with a strategy to minimise overall movement in a session, which would correspond to staying at the last rewarded port until stimulus onset, but instead suggests mice optimized for fast travel time toward the high reward patch once revealed. Strikingly, in a control condition where mice chose between two options of equal value, they developed distinct decision preferences and start position patterns that matched the optimal solution (Fig. S4). These findings highlight the importance of studying decision-making in settings that allow naturalistic behavioural variance.

Our task differs from previous rodent competition paradigms in several key ways ^49–51^. First, animals chose between two distinct options, and the reward-maximising strategy sometimes favoured the low-value patch. This enabled us to quantify how animals flexibly shifted their decision preference under competitive pressure (Fig. 2b,c). Second, the reward locations were unpredictable across trials, discouraging spatial monopolisation to ‘guard’ sources. As a result, dominant animals may have had less physical advantage in our game than in tasks with stable reward zones ^49,50^. We found that mice adjusted their strategies more when facing faster opponents (Fig. 2g), which may link to prior findings that subordinate animals adapt more than dominant animals during social interaction ^52^. Third, because mice actively surveyed their environment before each trial (Fig. 1g), their relative distances to reward patches and thus the corresponding reward probabilities varied from trial to trial. This notably increased the complexity of the decision process. Remarkably, mice integrated the instantaneous positions of both ‘self’ and ‘other’ to make rapid and highly flexible choices on each trial (Fig. 2f, Fig. 3c).

To benchmark these strategies, we used multi-agent RL simulations ^53^. ‘Social’ DQN agents successfully reproduced the opponent sensitivity observed in mice, whereas ‘socially blind’ agents could not, even when allowed various timescales of learning (Fig. 3). With noiseless information, the ‘social’ agents were more strongly influenced by opponent position than mice. It would be valuable to test whether agents with noisy, biologically plausible sensory inputs would exhibit behaviour more closely matching animal data. Conversely, some context-specific preference shifts could be mimicked by learning alone (Fig. S8, Fig. S9), illustrating the importance of using artificial agents to generate synthetic benchmarks, especially in tasks where key variables are outside of experimental control. Indeed, initial analysis of ‘socially-blind’ DQN agent behaviour suggested spurious sensitivity to opponent position: an impossibility by design. This artefact arose from analysing only the winner’s decisions, because the probability of winning depended on the position of opponent (Tab. S23). Identifying this issue in synthetic data led us to reanalyse the mouse data to include the inferred loser’s choices, thereby avoiding this potential pitfall.

The dynamical model provided further insight into the decision process and how it changed across contexts and trial configurations (Fig. 4). While bistable attractors have long been used to model decision-making ^41–43,54,55^, and dynamical systems have been used to describe aspects of social behaviour ^44,56–59^, we are not aware of previous work applying these models to behavioural data in competitive social foraging. Unlike the RL simulations, which required explicit discounting of reward, the attractor dynamics alone accounted for animals’ tendency to choose the low patch when nearby. The connection between delay-discounting and attractor dynamics remains underexplored (but see ^60^); it would be interesting to investigate under what regimes do attractor dynamics lead to changes of mind, e.g. ‘dynamic inconsistency’ in intertemporal choice, traditionally modelled with hyperbolic discounting ^61^.

The dynamical system approach also generated synthetic ‘neural’ data and made specific predictions about the time course of decodability for key task variables. The distinct inputs to the model, such as value input and opponent modulation, point to concrete neural candidates for investigation. For example, the nucleus accumbens and amygdala have been proposed to convey value signals to prefrontal cortex during social decision-making ^62^. Our study provides a quantitative foundation for testing such hypotheses. Currently, we model decisions as dynamics reaching a threshold followed by a ballistic movement. A future extension that aligns the dynamical system’s state with the animal’s continuous trajectory toward the chosen patch could better capture the embodied nature of these navigational foraging decisions ^63^.

The neuroscience community is increasingly interested ^64,65^ and equipped ^66–70^ to study innate and learned social behaviours ^71–76^. Here, we extend behavioural, analytical, and computational tools from solo decision-making to a competitive, multi-agent setting, offering a novel framework for uncovering mechanisms of flexible, value-based choices in a fast-changing and social environment.

## Methods

### Subjects

All experiments were performed under the UK Animals (Scientific Procedures) Act of 1986 (PPL PD867676F) after local ethical approval by the Sainsbury Wellcome Centre Animal Welfare Ethical Review Body. Ten adult C57BL/6J mice (8 males and 2 females; Charles River Laboratories), at least 8 weeks old at the start of the experiments, were used in this study. Mice were co-housed with their litter mates, with ambient temperature and humidity, and 12-h reversed day-night cycles. Mice were water-restricted before the start of behavioural training, with *ad libitum* access to food. On training days, mice received water as reward inside training rigs. Supplementary water was provided for mice to maintain a stable body weight (around 90%, but at least 85% of their starting body weight).

### Behavioural apparatus

#### Hardware

All experiments were conducted in custom-designed and manufactured open arenas (Fig. S1; NeuroGEARS Ltd., UK; FabLabs, SWC). Each arena consisted of an elevated base and 8 white acrylic wall panels, forming an octagonal space 80 cm in diameter and 50 cm in height. To allow visual stimuli presentation on the arena base ^77^, a short throw projector (HD29HST, Optoma) was used to project images onto a tilted mirror on the room floor, which then reflected the visual stimuli onto the arena base. The arena base was made from a transparent acrylic sheet with a rear projection film (Pro Diffusion, Pro Display) glued underneath, located 70 cm above the floor. Calibration was conducted using custom software in Bonsai ^67^ to ensure approximate equal luminance across the arena base and sufficient dynamic range for visual stimuli presentation. Each arena wall panel contained two nose-poke ports (OEPS-7246, Open Ephys), 3 cm above the arena base, for nose-poke detection upon infrared (IR) beam breaking. Each port also had an LED for additional visual aid; as well as a water spout, which was connected to a water reservoir via a computer-controlled pinch valve (161P011, NResearch) for reward delivery. To facilitate real-time mouse position tracking in the dark, custom made infrared (IR) illumination strips (850 nm LED) were mounted above, around, and below the arena. Videos were captured by an IR camera (BFS-U3-16S2M-CS, Flir; LN048 lenses) at 50 fps, mounted 124 cm above the centre of the arena base. A second monochrome camera with a bandpass filter (2020OFS-585, OFS) was used to monitor visual stimuli and animals’ ongoing behaviours.

#### Data acquisition and task control

Custom written software in Bonsai ^67^ was used to read and control all sensors and effectors in the arena via Harp devices (OEPS-7240, Open Ephys; Fig. S1a). In brief, each arena wall’s nose-poke ports and water valves were controlled by one Harp device (8 devices in total); and all 8 devices were synchronised using 2 clock Harp devices (OEPS-7231, Open Ephys). Projector visual stimuli presentation and camera video acquisition were triggered and synchronized by Bonsai. Behavioural task logic was written in Python-based software; and an Application Programming Interface (API) was developed to communicate between Python and Bonsai using a messaging library (https://github.com/glopesdev/pyOSC3). Data acquisition, video synchronization, data storage and access format were based on an API developed by the Aeon Foraging Behaviour Working Group ^78^.

### Behaviour

#### Solo task

In the final stage of the solo task, mice (n = 8) were guided by visual stimuli to perform value-based decision-making in the Octagon arena. The start of each trial was cued by a background colour change in the arena floor from black to gray. After a variable delay (0.5–1.5 s), two floor-projected visual stimuli (grating or white light triangular slices) were presented to indicate the location and value of the active reward patches for that trial. One visual pattern was associated with a high reward (3–5 units of 2.5 µL water reward, *p* = 1), and the other pattern was associated with a low reward (1 unit reward, *p* = 0.3–0.6). The luminance of the two visual stimuli were balanced to avoid perceptual biases; and the mapping between visual stimuli to reward sizes were counter-balanced across mice. The stimuli remained on until mice indicated their choice by nose-poking into one of the available patches within a response window (30 s from stimulus onset), which in turn triggered the corresponding reward. Poking into either of the two nose-poke ports on a single wall resulted in identical outcomes. Failure to respond within the 30 s response window was considered a ‘miss’ trial. Response time (RT) was defined as the time between stimulus onset and animals’ choice; and only trials with RTs shorter than 15 s were included in further analyses (mean RT: 1.77 s). All trials ended with an intertrial interval (ITI) of 6–12 *s*. Mice were not constrained to be at a particular location in the arena before each trial start. To increase the probability of mice attending to visual stimuli on the next trial, a new trial would only start after a minimum of 2 s where mice were not poking into any of the 8 walls. In each session, 80% of trials were ‘choice trials’ where mice chose between two different reward options; and 20% were forced trials where mice chose between two identical reward options in different locations. The locations of patches were randomized across trials, and the two patches could be 45^◦^ apart (neighbouring), 90^◦^ apart, or 135^◦^ apart. Sessions ended when animals showed increased miss rate as a sign for satiation. A control group of mice (n = 2) were trained to associate the grating and light floor stimuli with equal reward sizes (3 units); and their solo preferences for different stimuli were characterized and compared to the experimental group.

#### Training procedure

In the first behavioural session, mice were habituated with the arena and learned to poke into LED-illuminated ports for water reward. To aid operant conditioning in this large arena, all 16 ports were illuminated on each trial to start with and poking into any of the ports within a 60 s response window would trigger a reward (3 units reward). The number of available ports on each trial gradually reduced upon every 10 successful reward collection, until only two ports from 1 randomly chosen wall were illuminated on each trial. Mice typically learned to follow LED-illuminated ports in the Octagon arena within the first session. In the next 1–2 sessions, mice were trained to follow floor-projected visual stimuli rather than port LEDs for reward. On each trial, a single triangular stimulus (grating or white light slice) would appear on the arena floor to cue a specific wall as the active reward patch for that trial. After a variable delay (7–15 s), if no response in the active patch was registered, the corresponding port LEDs would light up to facilitate transfer learning. When mice could reliably respond to floor stimuli without port LEDs, only floor stimuli would be presented on each trial, and a response within a 30 s window would trigger the corresponding reward for that visual stimulus. To reduce anxiety, cage mates were trained together in these initial shaping steps, after which mice started solo training to discriminate visual stimuli associated with different reward sizes. As described above, two floor stimuli were presented on each trial and mice needed to choose one of the two active patches for reward. Poking into any non-active patch (extremely rare) was recorded but not punished or rewarded. Animals with stable preference for the high reward patch (*>* 70–80%) then moved on to competitive social sessions.

#### Competitive social task

The trial structure in the competitive task was identical to the solo task. Two well-trained mice (cage mates) were placed in the arena together. On each trial, only the first mouse to report its choice would be rewarded for their choice; and the slower mouse would receive no reward. Similar to the solo task, new trials would only start after a minimum of 2 s where neither of the two mice was poking into any of the 8 walls. On each trial, the winner mouse that received the reward was referred to as ‘self’ and the slower mouse was referred to as ‘other’ or the opponent; and ‘self’ identification varied across trials. Only cage mates with the same sensory-reward mappings were paired together for competition. For each pair of competing mice (n = 8 pairs), we conducted 3–8 triplets of solo-social-solo experiments. Social sessions lasted longer than solo sessions to approximately match the number of rewarded trials per mouse (social 118 versus solo 151 trials/mouse/session).

### Video tracking and identification

To track mouse identification during multi-animal sessions, we bleached back fur in one of the two competing mice so they could be easily distinguishable in real time by Bonsai tracking algorithms and post-hoc by SLEAP ^66^. Briefly, mice were anesthetized with isoflurane (2%). Human hair bleach was mixed according to the manufacturer’s instructions; and applied only to the top of animals’ fur to avoid skin irritation. The bleach stayed on for 7 min before it was wiped off and rinsed with saline. Mice were then placed back to a heated cage to recover from anesthesia. Fur bleaching was always conducted before solo sessions to ensure that there was no change in baseline task performance after brief anesthesia or the bleaching procedure. Bleaching was repeated when needed, usually after 2-4 weeks.

The infrared (IR) camera was used to track animals’ positions and identifications in real time at 50 fps. Briefly, the IR camera imaging frames were thresholded to expose the region of interest (ROI) corresponding to the mouse (the black area of the mouse’s fur versus the white arena floor). In social competition experiments, the bleached fur reduced the ROI area of the bleached mouse, allowing the two mice to be distinguished based on the ROI size (Fig. S1d). On each frame, the central position (centroid) and area size of a large ROI (non-bleached mouse) and a small ROI (bleached mouse) were tracked and recorded. We used the centroid positions and area sizes at the time of response to determine which mouse was the winner (‘self’) on each trial. Tracked body size could vary depending on animals’ varied postures within a session. To ensure correct identification using our method, we plotted the distributions of tracked body sizes across all the frames in each pair of solo control sessions. If the two distributions were non-overlapping, we proceeded with the social experiment on the next day; otherwise bleaching was repeated. Identification tracking was further validated post-hoc using SLEAP ^66^.

After extracting animals’ positions from the custom Bonsai pipeline, the trajectories underwent further offline processing, to fill any gaps in the positional data using linear interpolation, and to smooth outlier data due to tracking artifacts. Although the loser’s choice was not rewarded, a subset of analyses required us to include both the winner’s and loser’s choices to eliminate spurious correlations. We analyzed post-processed trajectory data and inferred the loser’s choice by estimating which of the two active walls they were closer to, 1 second after the winner’s choice. We only included trials where the loser’s distance to their chosen wall was less than 12 cm, so trials where they were at a different response port (e.g. still at a reward port from a previous trial) were excluded. Using a stricter criterion did not change any of our main conclusions.

### Behavioural data analysis

#### Solo behaviours

Analyses of behavioural data were conducted using custom software written in MATLAB (Mathworks, 2021b) and Julia 1.10^79^. For all analyses, we used a coordinate system where the centre of the octagon was [0,0] and the distance to the midpoint of the patch walls was 1. To characterize post-learning performance (Fig. 1), we included the last 30 days of stable solo training from each mouse. Generalization of value preference across patch locations (Fig. 1f) was demonstrated by concatenating all the choices from 8 mice, and plotting the probability of choosing the high patch, P(high), as a function of actual high wall locations. To quantify response time (RT) differences across trial types (Fig. S2), we calculated the median RT for each trial type per mouse, and plotted the log ratio of RTs, normalized to trials where mice were ‘forced’ to choose between two high patches. Within-subject bootstrapped tests were used for significance testing. The relationship between forced trial log RT ratio and choice trial value discrimination was quantified using linear correlation across sessions (Fig. S2c). To test whether the high/low mice were statistically closer to the arena centre at trial start compared to the control mice, we used the Receiver operating characteristic (ROC) analysis and estimated the area under the ROC curve (AUC) between distributions of normalized distances to the arena centre on concatenated trials across mice groups.

We quantified the spatial effects on animals’ value preference in two ways. First, for visualisation, we rotated and reflected varied trial configurations so that the high reward patch was at direct north, and the low reward patch was east to the high patch (Fig. 1g,h). Trials were then binned based on start positions relative to the high and low patches, and averaged P(high) was plotted for each start position bin in the Octagon space (Fig. 1j). Second, for statistics and model comparisons, we calculated the distance from the mouse’s start position to the high patch and to the low patch on each trial, and then used distances rather than positions to evaluate spatial effects on performance (Fig. 1k). The first approach was visually salient, whereas the second approach allowed us to combine trial conditions where the two patches were 45^◦^ or 90^◦^ apart.

To assess animals’ overall solo strategies, we fit generalized linear mixed models (GLMM, implemented in MATLAB function fitglme) where animals’ choice on each trial was a logistic function of intercept and their distances to the high and low patches at stimulus onset as fixed effects, as well as within-subject random effects (See Statistical Appendix for details). We compared 4 models: model 3 (Tab. S3) includes distance to low only; model 4 (Tab. S4) includes distance to high only; model 2 (Tab. S2) includes both distances as regressors; and model 1 (Tab. S1) includes both distances and their interaction (Fig. S3c). Twenty-fold cross-validation was used to calculate per trial normalized log likelihood for model comparisons (Fig. S3e), where *P* values were calculated using paired bootstrapped tests across folds (Tab. S5). Significant effects of distances and their interaction on P(high) were quantified based on regressor coefficients and nested model comparisons. We also fit model 2 to each animal separately and reported the regression coefficients per animal (Fig. S3d). All regression tables are shared in the Statistical Appendix.

#### Competitive behaviours

To assess how competition shifted animals’ overall decision preference, we calculated P(high) on winning choices in each social session, and compared that to P(high) on the corresponding solo sessions one day before and one day after (Fig. 2c). For each pair of animals, trials from both animals’ solo sessions were concatenated to calculate the average solo P(high) for that day. Paired *t*-tests were used for statistical testing across 38 session triplets. Further separating trials from each social session into the first and second halves did not reveal any consistent change due to slow adaptations within the social sessions (Fig. S5d). To ensure that the preference shift observed was not merely a speed-accuracy tradeoff, we conducted a control analysis that simulated selection of the faster choice between each pair of mice without the element of competition. Specifically, for each trial in the social session, a pair of trials were randomly chosen, with replacement, from two animals’ solo control sessions 1 day before; and only the faster trial from each pair was included into an RT-corrected solo control session (Fig. S5a-b). We found similar results using this conservative control analysis.

To visualize how animals’ value preference shifted under competition as a function of self positions relative to goals, we used similar methods as described for Fig. 1j,k, but considering whether the trials came from solo vs. social sessions (Fig. 2d-f). For statistics, we fit GLMMs on concatenated solo and social trials, where animals’ choice on each trial was a logistic function of the chooser’s distances to the high and low patches at stimulus onset as well as whether the trial was from a solo or social session (social context) as fixed effects, and within-subject-pair random effects (See Statistical Appendix for details). We compared 3 models: model 3 (Tab. S8) only included self distances to goals and their interaction as regressors (Fig. S6d); model 2 (Tab. S7) included self distances to goals and the social context (Fig. S6c); and model 1 (Tab. S6) included additional interactions between self distances and the social context (Fig. S6b). Twenty-fold cross-validation was used to calculate per trial normalized log likelihood for model comparisons (Fig. S6f), where *P* values were calculated using paired boostrapped tests across folds (Tab. S9). Significant effects were quantified based on regressor coefficients and nested model comparisons. We also fit model 2 to each animal separately and reported the regression coefficients per animal (Fig. S6e).

To test whether mice shifted their strategies differentially when playing against varied opponents, we extracted choices by each mouse in the social session, and compared how P(high) changed from their respective solo performance the day before as a function of opponent features. Specifically, we estimated opponent speed (1/response time) and opponent preference for the high reward patch on solo days, and correlated these opponent features with how much self preference was shifted when competing against these opponents using linear correlation. We found that animals shifted their strategy more against fast opponents (Fig. 2g). To ensure this correlation did not result from analyzing winners’ choices only, we included inferred losers’ choices and still found a significant correlation between opponent speed and self strategy shift in social sessions (*r* = .24*, P* = .04*, n* = 70 sessions).

Finally, we characterized how opponents’ instantaneous positions relative to goals modulated animals’ choices on a trial-by-trial basis (Fig. 3a-d). To avoid any spurious correlation between the probability of self winning and opponent’s positions across trials (Tab. S23), it was essential that we included both the winners’ choices and the inferred losers’ choices for this analysis. For visualisation, we concatenated all social trials and separately plotted two conditions across all self positions: when the opponent was far from the high patch (*>* 60 percentiles of all opponent distances; Fig. 3a) and when the opponent was close to the high patch ( *<* 40 percentiles; Fig. 3b). We also plotted the mean difference for each self position bin (Fig. 3c). For statistics, we used GLMMs (using MixedModels.jl ^80^) to model each social trial as a logistic function of the chooser’s distances to the high and low patches at stimulus onset as well as the opponent’s distance to the high patch as fixed effects, and within-subject random effects (See Statistical Appendix for details). We compared 3 models: model 1 (Tab. S10) only includes self distances to goals and their interaction as regressors; model 2 (Tab. S11) includes self distances to goals and the main effect of the opponent distance to the high patch; and model 3 (Tab. S12) includes additional interactions between self distances and the opponent distance to the high patch. Significant effects were quantified based on regressor coefficients and nested model comparisons (likelihood ratio test, Tab. S13). The best fit model (model 3) was visualized in Fig. S6g,h.

### Simulations of optimal start positions

To determine optimal start positions (Fig. 1i), we assumed that agents aim to minimize the time (and therefore distance) from the start position to the desired patch. We performed these simulations in Julia (1.10) using the Meshes.jl package to define an octagon with an inradius (distance from the centre to the midpoint of each edge) of 1. Then, we created a grid of start position points (200 *×* 200) uniformly spaced from [-1,1] and determined the points that were inside the octagon. Of the 40000 points, 32756 were inside the octagon. We found that this number of start position simulations generated a smooth visualisation ^81^.

For high/low mice to maximize water intake in solo sessions, animals almost always chose the high reward patch. In order to minimize the distance to the high patch across trials, the best start position would be the one that had the shortest average distance to a single randomly appearing wall, so we computed the mean distance from every point in the octagon to a randomly appearing wall (Fig. 1i). If a mouse instead wanted to choose the closer one of the two available walls, as in ‘equal’ mice, then the optimal start position is the position in the octagon with the shortest average distance to either of the two randomly appearing walls (for both 45^◦^ and 90^◦^ trials; Fig. S4h,i).

### Reinforcement Learning: DQN model

The reinforcement learning (RL) agent employed Q-learning, a form of off-policy temporal difference control algorithm that learned to estimate the action value function through iterative updates. In general, the goal of RL is to learn a policy, *π*, that maximizes the expected cumulative reward over time. Following the approach of Mnih et al. ^40^, we implemented a Deep Q Network (DQN) to approximate the optimal action-value function:

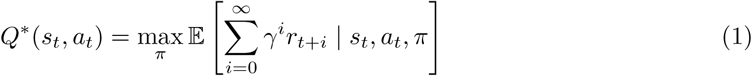

where *Q*^∗^ is the maximum sum of all the future rewards *r_t_*, discounted by *γ* at each time step *t*, if the agent follows the policy *π*.

Our environment was a 64 *×* 64 pixel image (Fig. S7a), providing a bird’s-eye view of the Octagon arena with positions of the reward patches and agent positions. The agents were depicted as simple geometric shapes: circles and squares. The action space consisted of the 8 possible wall choices in the arena, along with a null action (i.e., an action which avoids action costs but gets no reward). Rewards, *R* for each agent depended on the value of the chosen patch. Additionally, the reward was discounted by *γ^d^* and an action cost, *AC*, was subtracted from the reward, representing the effort required for the agent to traverse the Octagon arena and reach the selected wall. This action cost was calculated under the assumption that the agent followed a direct path from its starting position to its chosen destination, which is why the cost term is written as the sum of a geometric series. The reward for a specific choice was expressed by the following equation:

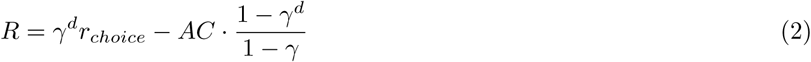

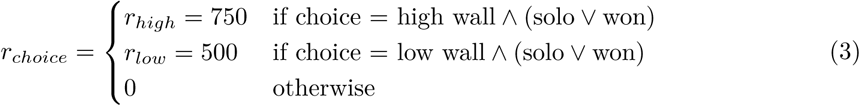

where *d* was the distance of the agent from it’s starting position to the wall chosen (measured in pixels), *γ* = .98*, AC* = 5. In the social context, the winner was determined purely based on the distance of the agents to their choice. The parameters values (*r_low_, r_high_, γ, AC*) were chosen to qualitatively fit the mouse behaviour in the solo context, other hyperparameters were described in Tab. 1. Each episode consisted of a single transition (i.e., a single action to choose the wall), and decisions were modeled solely on the initial positions of the agents at trial start. An ‘episode’ for the DQN corresponded to a single trial experienced by mice in the experimental arena.

**Tab. 1:** DQN Hyperparameters.

| Hyperparameter | Value |
| --- | --- |
| Learning rate ($\alpha$) | 0.0001 |
| Softmax temperature ($\tau$) | 5 |
| Batch size | 128 |
| Replay buffer size | 1000 |
| Recency Weight ($rw$) | 0.1, 0.01, 0.001 |
| Loss function | SmoothL1Loss |
| Optimizer | AdamW |

The DQN took the image input of the environment and had 2 convolutional layers followed by 3 fully connected layers. The final layer of the DQN represented the Q-value for every available action (9 in total) in that state for the agent, which was used to choose an action by applying a softmax with temperature (*τ*). The update step of the DQN involved sampling from a ‘replay buffer’ of past experiences (1,000 episodes). Episodes were sampled using a geometric distribution *G*(*p* = *rw*) to prioritize recent observations, where *rw* is the recency weight. When *rw* was small, the buffer was sampled more evenly, and as *rw* got larger, more recent experiences were oversampled (Fig. S8a).

#### Simulation Details

In order to facilitate comparison between mice and DQN, we first pre-trained the CNN layers of the DQN to label trials as solo or social trials (details see below). We reasoned that mice know whether or not another mouse is in the arena, so we didn’t want this aspect of learning to slow down task training. Likewise, we first trained the DQN in the solo context until their behaviour was qualitatively similar to expert mice (5000 episodes, which is similar to the number of trials in 30 sessions for a mouse). Then we alternated 500 episode blocks of solo and social contexts. Since the DQN needed to distinguish ‘self’ and ‘other’ from the bird’s eye-view input, 10 DQN learned that self was a circle and 10 DQN learned that self was a square. To mimic the individual differences in mice, we trained DQN using 10 random seeds to initialize ^82^ the dense layers for each agent type (circle and square), yielding 100 pairwise simulations (10 circle vs. 10 square agents). These simulations were repeated (with the same seeds) for social vs. socially blind agents and for three levels of recency weighting (rw; Fig. S8), giving 100 *×* 2 *×* 3 = 600 simulations. On each trial, the agents were randomly placed within the arena, ensuring that the glyphs representing the agents in the environment were far apart to avoid any occlusion. The position of the low and high walls was also randomly chosen throughout each episode, maintaining the relative positions between them as in the experiments (45^◦^ and 90^◦^ separations) giving 32 possible configurations of the high/low walls. All the simulations and the DQN training was done in Python using PyTorch.

#### Pre-training

Before task-specific learning, the CNN was trained on a binary classification task designed to differentiate between solo and social contexts based on the input image. In the solo context, only one agent (either a circle or a square) was present, whereas in the social context, both agents were included in the image presented to the network. This pre-training enabled the network to differentiate between solo and social contexts without acquiring strategies specific to the decision-making task. Only the CNN layers from the pre-training were used as initialization in the DQN, allowing the model to continue learning and adapt these parameters during the task-learning phase.

#### DQN Analysis

To plot P(high) for the DQN as a function of position in the solo context (Fig. S7d), we took the last 2000 trials from the first solo block for the 20 DQN agents and averaged across agents. For the plots of DQN behaviour in the social context (Fig. 3e-l), we took data only from the last social session (n = 500 trials/simulation *×* 100 simulations = 50 000 trials). For comparisons across the solo and social contexts, only the solo sessions before and after the last social session were included.

### Attractor modelling

We simulated a 2-node dynamical system to understand the contributions of a small number of interpretable factors to the decision processes in the Octagon task and to quantify how those factors changed between the solo and social contexts. There was no learning in these simulations, so we did not simulate pairs of dynamical systems to determine a winner on each trial. Instead, we simulated each trial of social sessions twice: once from the perspective of the winner and once from the perspective of the loser (for the trials where we inferred the choice of the loser, as in Fig. 3c). We ran the dynamical simulations in Julia 1.10^79^ using the DifferentialEquations.jl package^?^ and visualized the simulations using Makie.jl ^81^. The starting point for the implementation was the two-node system fit to data from an accumulation of perceptual evidence decision making task ^54^. Here, the activity of the two nodes, *X*, were governed by the following equations, where *V_i_*was the value of each patch, *ψ* governed the influence of opponent position, *W_E_*was the strength of self-excitation, *W_I_* was the strength of mutual inhibition, *E* and *σ* were the mean and standard deviation of the external drive, and *z_i_* was distance of the opponent to each patch (during social sessions). The initial value of each node, *X_i_*(*t* = 0), depended on the distance of the mouse to the respective patch, *d_i_*:

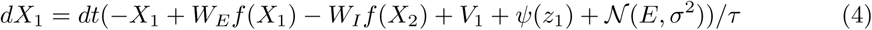

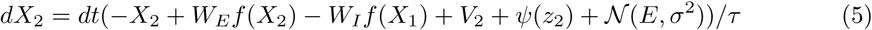

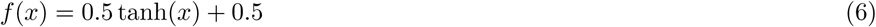

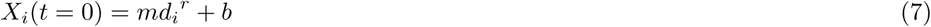

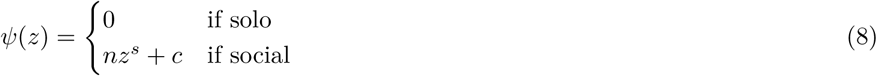

The solo model had 6 free parameters and the social model had an additional 3 free parameters (Tab. 2). Other parameter values were picked based on previous work Tab. 3^54^. We combined data across mice in order to find parameters that captured the overall patterns of behaviour. Initial attempts to find the maximum likelihood estimates of the parameters using standard minimization algorithms (Nelder-Mead) revealed that our system had many local minima. To overcome this, we randomly selected parameter sets (10 000 sets for solo and social) from a range and simulated each trial in the data 200 times to get a consistent estimate of the likelihood of the data given that parameter set. We took the best solo and social model from this ‘shotgun’ approach and then used minimization to find a nearby best model. The best models had a likelihood per trial of 0.6759 for solo trials (n = 24881) and 0.6388 (n = 10460) for social trials.

**Tab. 2:** Best model parameters.

| Parameter | Best Solo | Best Social |
| --- | --- | --- |
| $E$ | 1.239 | 1.916 |
| $\sigma$ | 3.775 | 3.899 |
| $V_H$ | 2.199 | 1.815 |
| $m$ | -1.155 | -1.623 |
| $r$ | 1.540 | 1.238 |
| $b$ | 1.471 | 0.777 |
| $n$ | n.a. | -0.741 |
| $s$ | n.a. | 1.000 |
| $c$ | n.a. | 1.534 |

**Tab. 3:** Fixed parameters for the dynamical models.

| Parameter | Value |
| --- | --- |
| $W_E$ | 1.5 |
| $W_I$ | 4 |
| $dt$ | 0.001 s |
| $\tau$ | 0.1 s |
| $V_L$ | 0.5 |

## Supporting information

Supplementary Video 1 Solo Examples

Supplementary Video 2 Social Examples

## Author contributions

C.A.D. and J.C.E. conceived the project, the arena and designed the behavioural task. J.C.E., B.C., and A.A. contributed to the software and hardware design and construction of the Octagon arena. C.A.D. and I.O. collected the behavioural data. C.A.D. analyzed the behavioural data, with J.C.E. and M.R’s inputs. Q.Y., M.R., and J.C.E. developed and analyzed the DQN simulations. J.C.E. performed the dynamical modelling. C.A.D. and J.C.E. wrote the manuscript, with contributions from M.R. and I.O.

## Acknowledgments

This work was funded by the Sainsbury Wellcome Centre Core Grant from the Gatsby Charitable Foundation (GAT3755) and Wellcome (219627/Z/19/Z); by a UK Research and Innovation grant (EP/Y008804/1); and by the Gatsby Initiative in Brain Development and Psychiatry (GAT3955). We thank members of the Duan laboratory and members of the Erlich laboratory for insightful discussions and advice; T. Branco, T. Margrie, and T. Mrsic-Flogel for comments on the manuscript; SWC Neurobiological Research Facility for animal support; and S. Townsend from FabLabs for contributions to arena hardware design and construction.

## Competing interests

The authors declare no competing interests.

## Code and Data Sharing

Code and data will be shared at the time of publication.

## Supplemental Videos

1. Supplemental Video 1. Solo Example Trials. https://shorturl.at/0nUYX
2. Supplemental Video 2. Social Example Trials. https://shorturl.at/R5C01

---

**Fig. S1:**
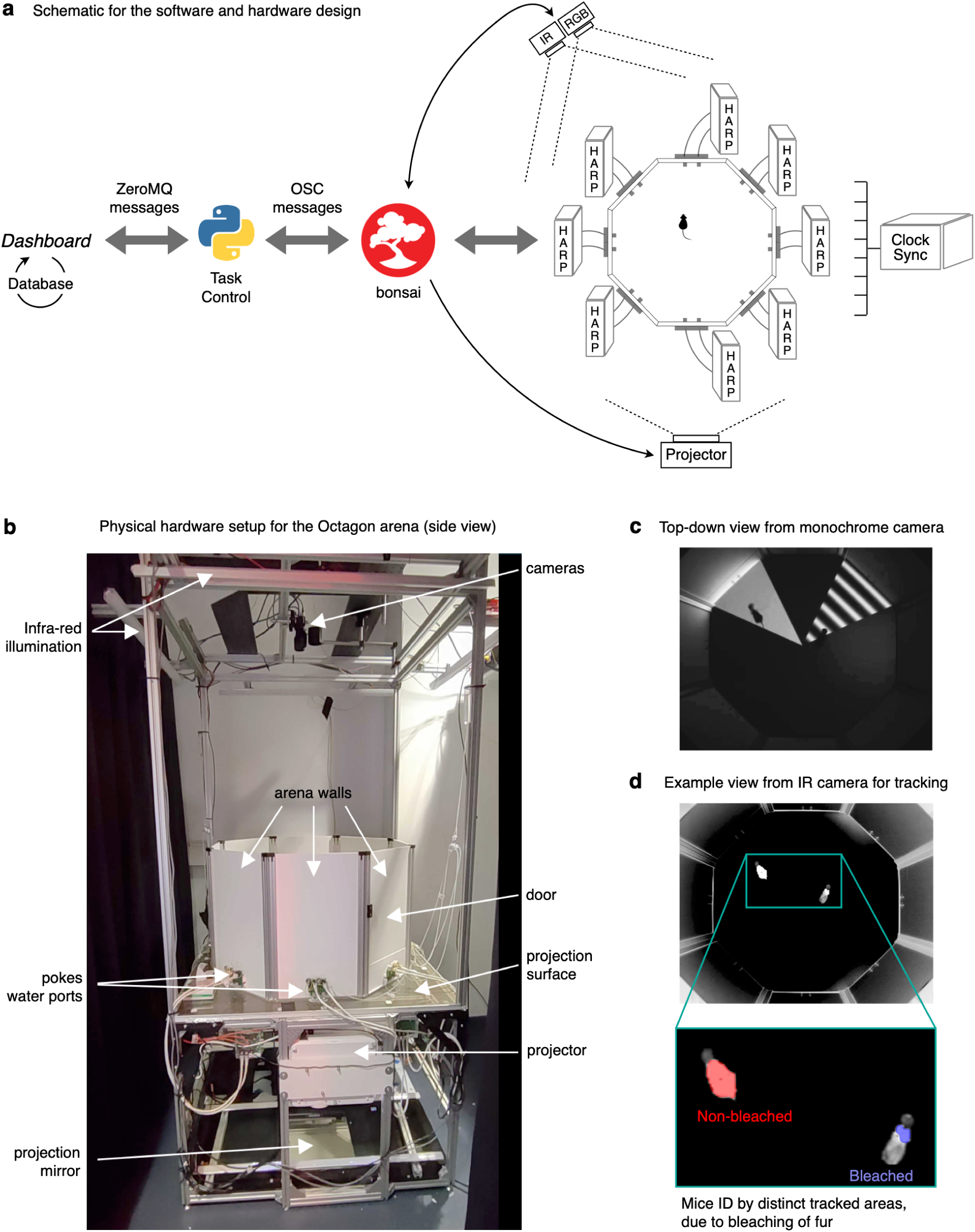
Design of the Octagon arena. **a**, Schematic for software (Bonsai, Python) and hardware (HARP) design, task control, data acquisition and storage. **b**, Side view of the Octagon setup. **c**, Example top-down view from the monochrome camera, where both the visual stimuli and mice could be monitored. **d**, Example view (same frame as **c**) from the IR camera, which was used for position and identification tracking. Note that the tracked area size for the non-bleached mouse was larger than that of the bleached mouse (Methods).

**Fig. S2:**
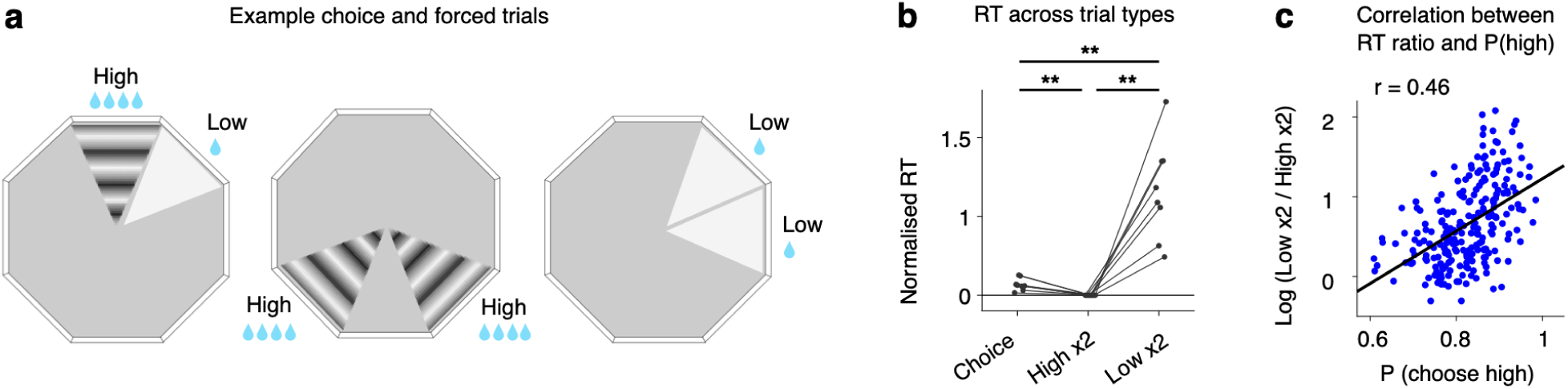
Forced choice RT and correlation with performance. **a**, Example choice and forced trials. **b**, Log ratio of response times (RT) compared to RTs on high-high forced trials, n = 8 mice. **, *P < .*01, paired bootstrapped tests. **c**, Session-wise correlation between forced trial RT log-ratio and choice trial performance, n = 235 sessions. r, Pearson’s correlation coefficient.

**Fig. S3:**
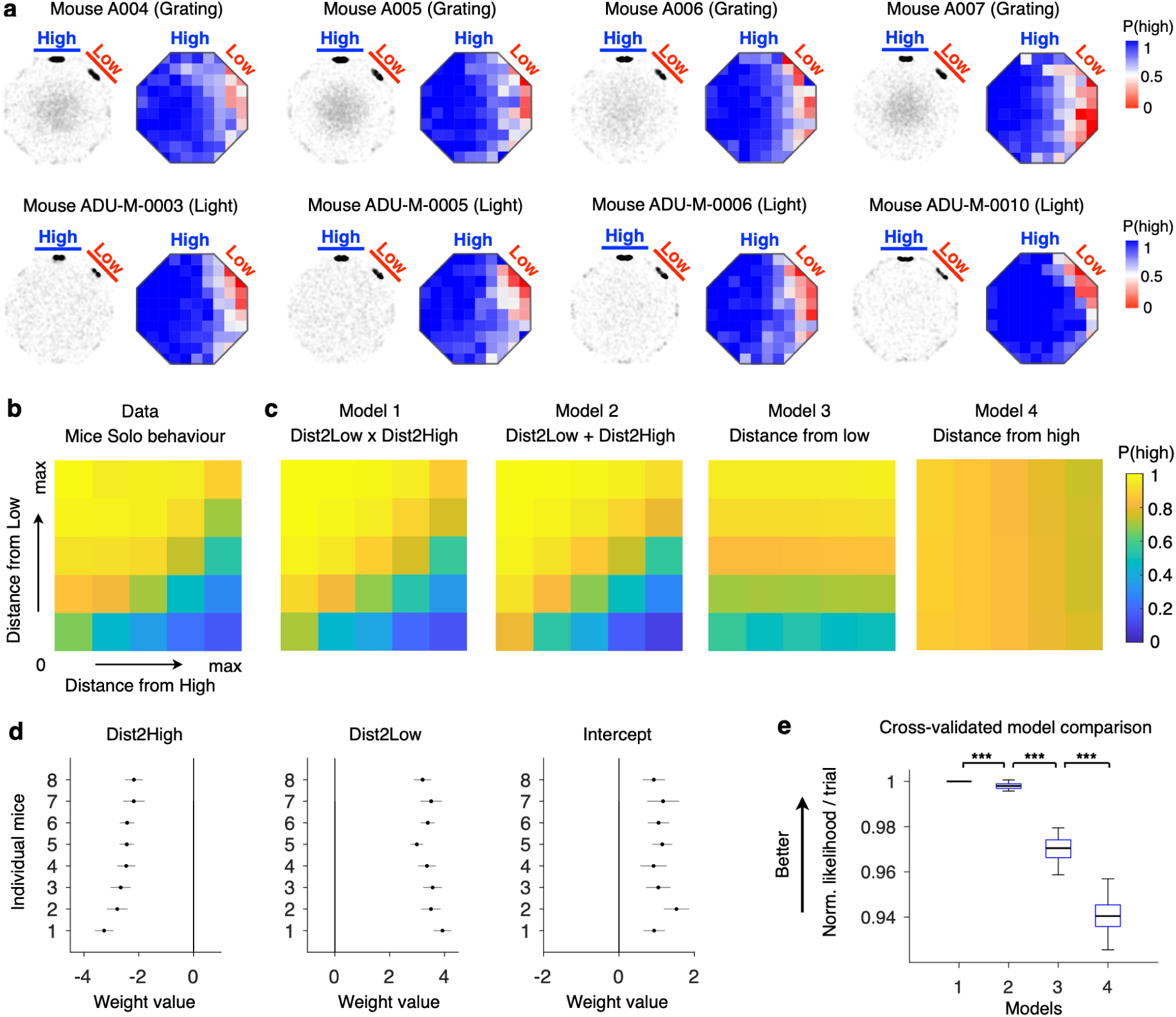
Solo preference for individual mice. **a**, Tracked positions at stimulus onset (gray) and at the time of response (black, left) and post-learning performance across varied start positions in the Octagon (right), rotated and reflected for visualisation, for 8 individual mice. All mice had a preference for the high reward stimuli, generalized across start positions and sensory-value mappings (top 4 mice, grating was associated with high reward; bottom 4 mice, light was associated with high reward). **b,c**, Performance as a function of normalized distance to the high reward patch and the low reward patch, for mice data (**b**, 29007 trials from 235 sessions and 8 mice), and for predictions from different GLMM models (**c**). Model 3 (Tab. S3) and model 4 (Tab. S4) only included the distance from the low patch regressor and the distance from the high patch regressor respectively; model 2 (Tab. S2) had both regressors as main effects; and model 1 (Tab. S1) had both regressors and their interaction. **d**, Regressor coefficients for the main effects of distance to the high patch; distance to the low patch; and the intercept from model 2, fit to each of the 8 individual mice. All mice had a general preference for the high patch, and their preference was modulated bidirectionally by the distances to the high and low patches. Error bars represent 99% confidence intervals. **e**, Normalized negative log likelihood per trial across different GLMM models, with 20 fold cross-validation. *** *P <* 0.001, paired bootstrapped tests. Including each distance regressor and their interaction significantly improved model performance (Tab. S5).

**Fig. S4:**
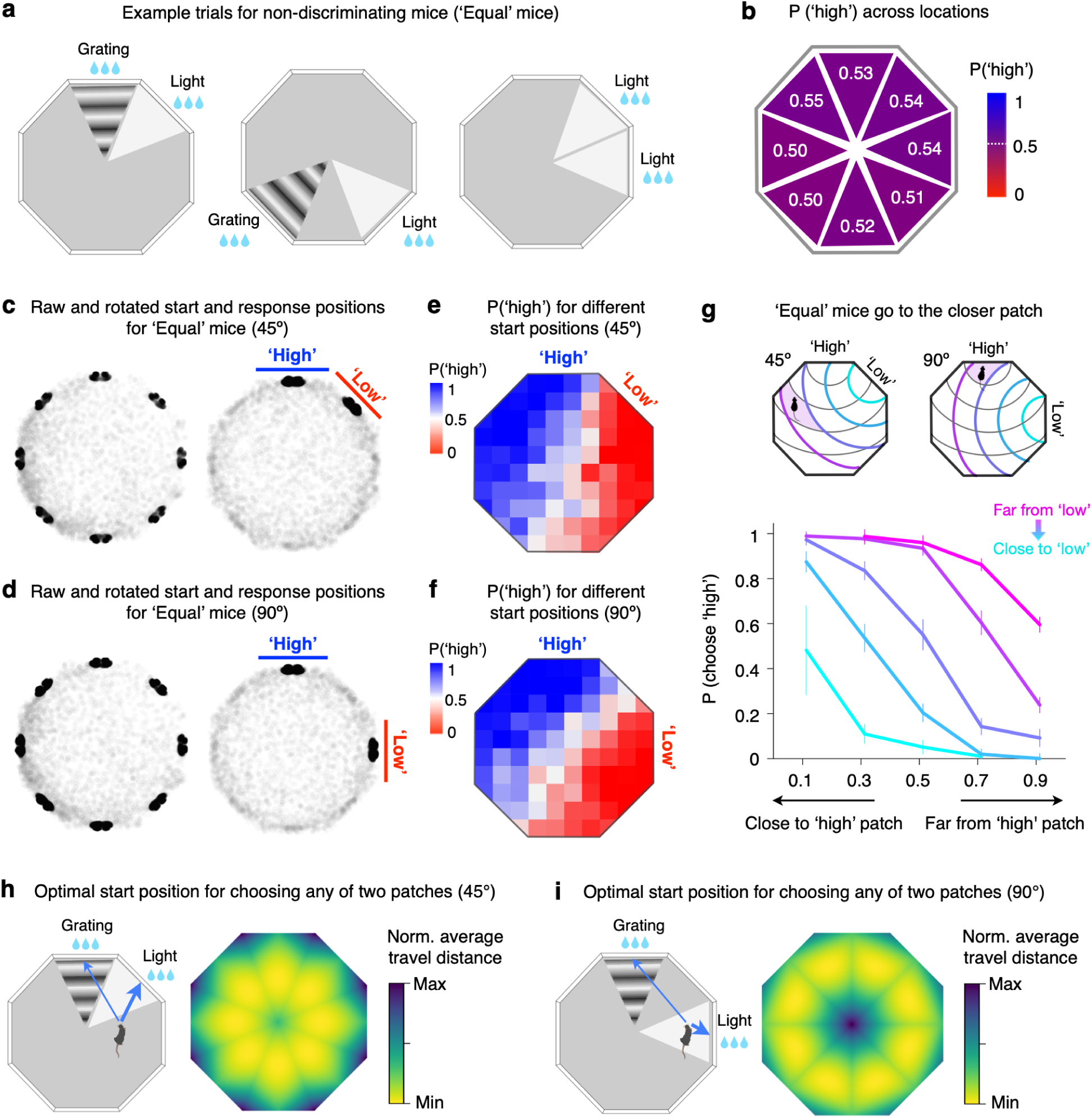
Behaviours in control (‘equal’) mice. **a**, Example visual stimuli for ‘equal mice’, identical to those used in the experimental group, except that each stimulus was associated with the same amount of reward (3 units). **b**, P(‘high’) is defined as the probability of choosing the light patch. ‘Equal’ mice did not develop a preference for any particular stimulus across different patch locations, in contrast to Fig. 1f. **c,d**, ‘Equal’ mice did not develop a strategy of clustering towards the arena centre at trial start, in contrast to Fig. 1g. **e-g**, ‘Equal’ mice made choices based on proximity, in contrast to the experimental group shown in Fig. 1j**,k**. **h,i**, Optimal start positions to minimize average travel distance if the goal is to choose the closer of the two randomly appearing patches, grating or light, when patches are 45 (**h**) or 90 (**i**) degrees apart. Note that start positions closer to edges can increase the probability of being close to one of the two patches, and is thus more optimal for a proximity-dominated strategy.

**Fig. S5:**
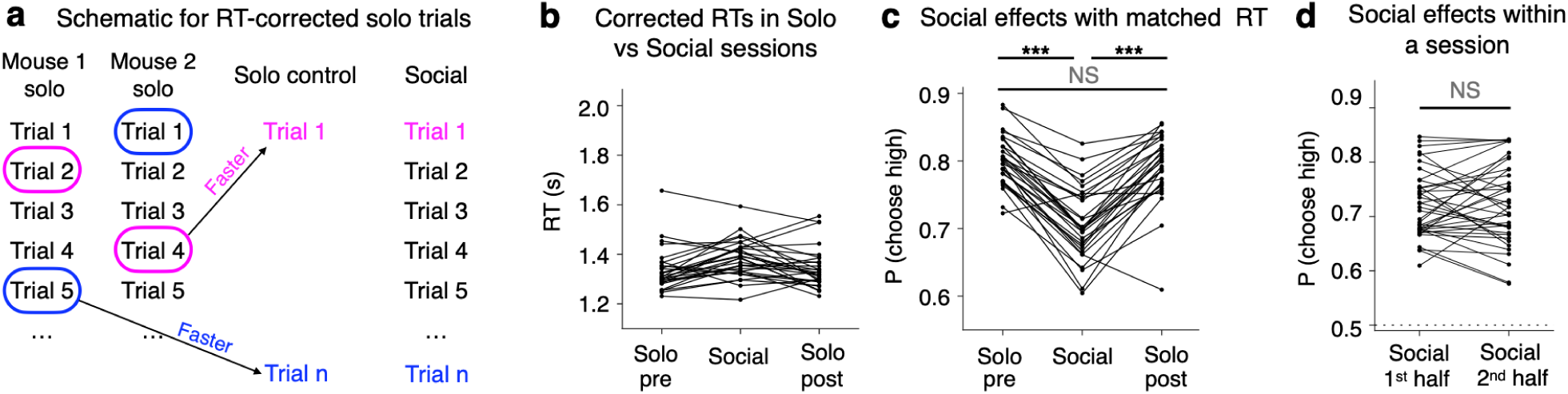
Preference changes across contexts when controlled for response time (RT) and session time. **a**, Schematic for how RT-corrected solo control trials were selected. For each trial in the social session, a pair of trials were randomly chosen, with replacement, from two animals’ solo control sessions 1 day before. To simulate race without competition, the faster trial from the pair was included as the corresponding solo control trial. **b**, Correction resulted in comparable RTs across contexts. **c**, With matched RT distributions, mice still decreased preference for the high reward patch under competition, similar to Fig. 2c. **d**, The first and second halves of social sessions showed comparable performance, arguing against a slowly changing preference due to non-social learning. ***, *P <* 0.001; NS, not significant, *P >* 0.05; paired *t*-tests.

**Fig. S6:**
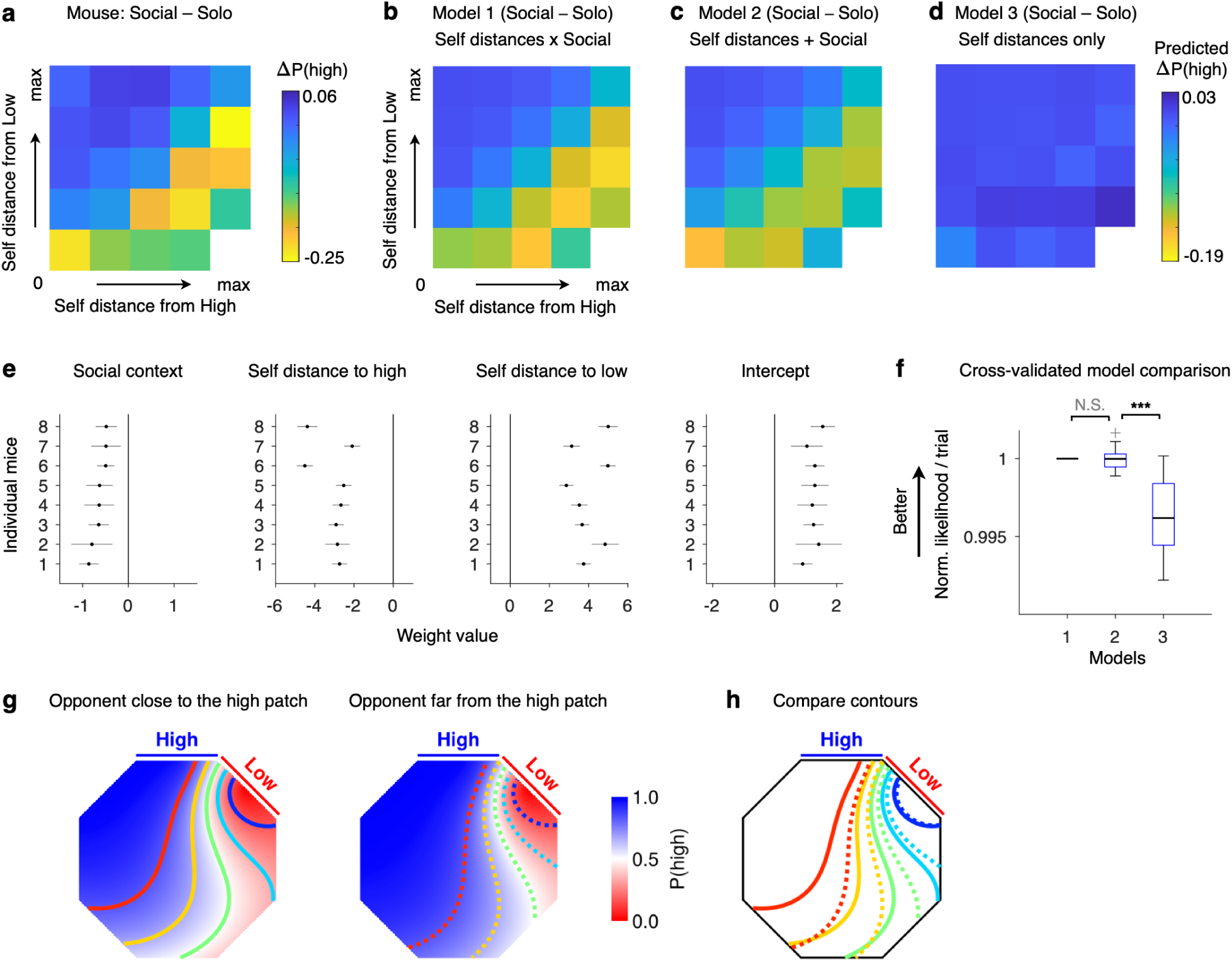
GLMM fits for the effects of social competition on performance. **a**, Decrease in the probability of choosing the high reward patch in social sessions (7035 trials, 8 mice) compared to solo sessions (9091 trials, 8 mice), as a function of the chooser’s distances to the high and low patches. **b-d**, Similar to **a**, for predictions from different GLMM models (Methods). Model 3 (Tab. S8) only included the distance regressors but not the social context; model 2 (Tab. S7) had both the distance regressors and the social context main effect; and model 1 (Tab. S6) included both distance regressors, social context main effect and their interactions. **e**, Regressor coefficients for the main effects of the social context; distance to the high patch; distance to the low patch; and the intercept, fit to each of the 8 individual mice. All mice had decreased preference for the high reward patch under social competition. Error bars represent 95% confidence intervals. **f**, Normalized negative log likelihood per trial across different GLMM models, with 20 fold cross-validation. ***, *P <* 0.001; NS, not significant *P* = 0.85, paired bootstrapped tests. Including the social context as a main effect significantly improved model performance. **g**, Visualisation of the GLMM fit to the mouse behaviour as a function of opponent distance to the high patch and self positions (Tab. S12). For the left panel other2high = min and for the right panel other2high = max. To aid comparison of the change in preference across the two panels, we drew contour lines at 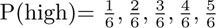 (in blue, cyan, green, yellow, red respectively), splitting the data into sextiles. To make this visualisation clearer, we only simulated 45*^◦^* trials. **h**, The contours from **g** replotted together for easier comparison. Note how the blue contour is relatively unchanged, and the difference between the contours grows larger as self is further away from the high and low reward patches, revealing the interaction effects of self and opponent positions on choices.

**Fig. S7:**
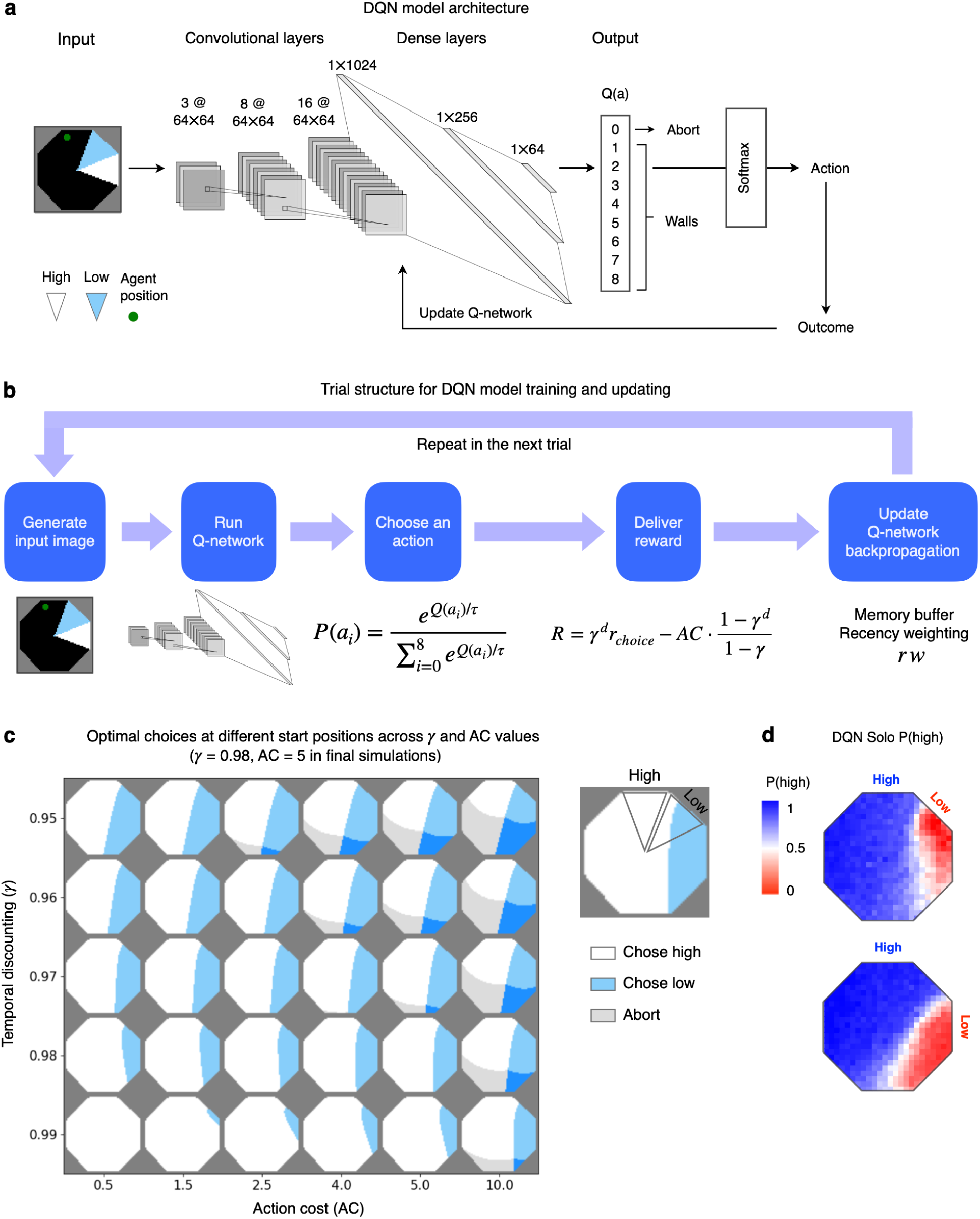
DQN details. **a**, Detailed model architecture. **b**, Detailed trial structure for input, choice generation, discounted reward delivery and network updating (Methods). **c**, Simulations for optimal choices at different start positions in the Octagon with parametrically varied values of temporal discounting (*γ*) and action cost (AC). *γ* = 0.98 and AC = 5 were chosen to qualitatively match mice solo preference. **d**, DQN performance on solo sessions, averaged across 20 agents, rw = 0.1, similar to mice solo performance in Fig. 1j.

**Fig. S8:**
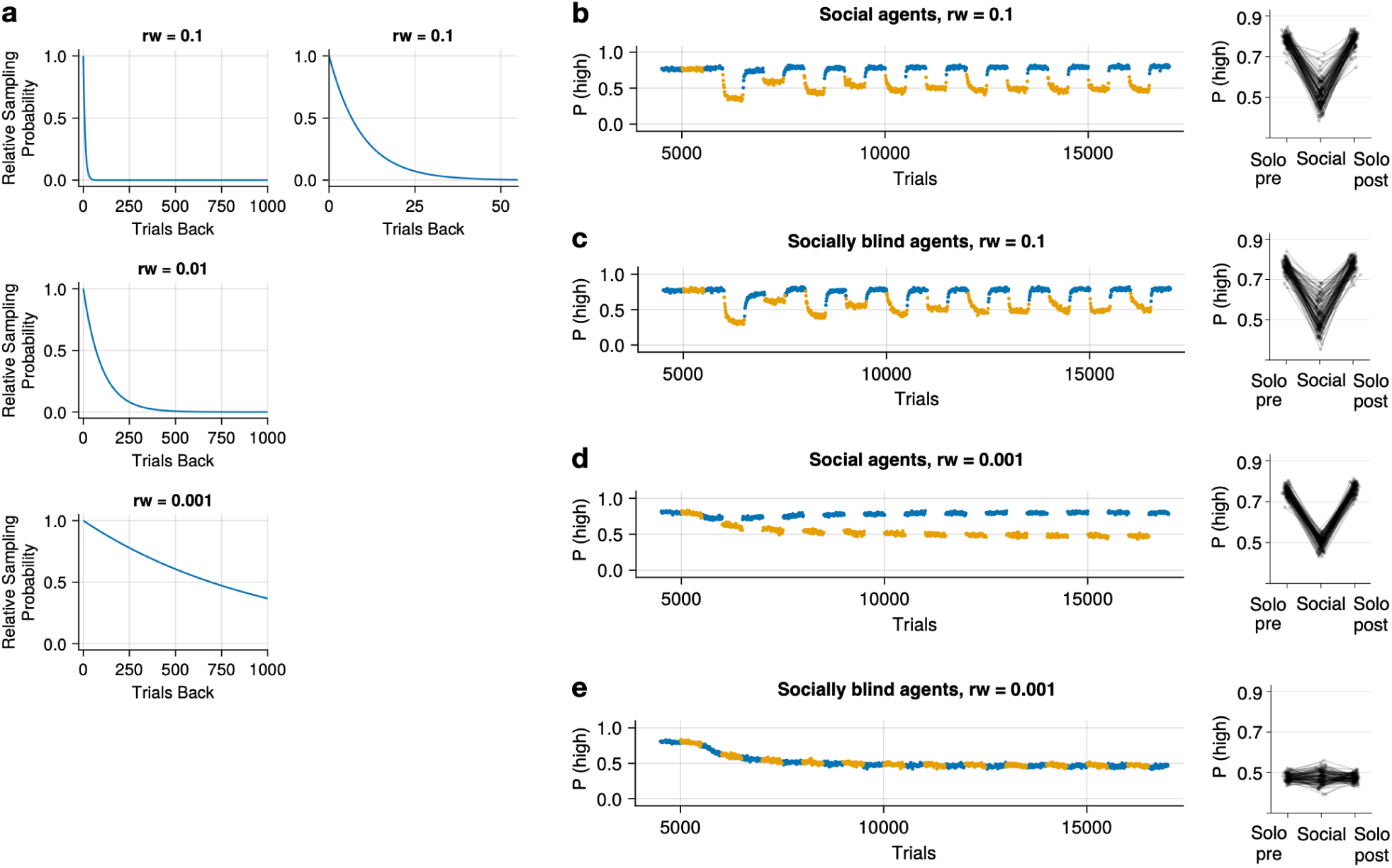
Influence of recency weighting on DQN behaviour. **a**, Visualisation of the priority sampling for different levels of recency weighting (rw). Each row shows the sampling for a smaller rw. For rw=0.1, we show a zoomed in plot (top right) to show that samples are from very recent history (e.g. *<* 50 trials back). **b,c**, Rapid learning (rw = 0.1) in ‘social’ and ‘socially-blind’ agents. Left, each point is the average P(high) of 10 trials, averaged across all simulations. Trials from solo sessions are in blue and social sessions are in orange (25 out of 60 training blocks are shown). Right, change in the probability of choosing the high patch across solo and competitive contexts, data from the last 3 blocks. For recency weighting (rw) of 0.1, both ‘social’ and ‘socially-blind’ agents switched strategies across blocks, but showed evidence of within-block learning. **d,e**, Similar to **b,c**, for rw=0.001. ‘Social’ agents had context-specific strategies, while the ‘socially-blind’ agents did not.

**Fig. S9:**
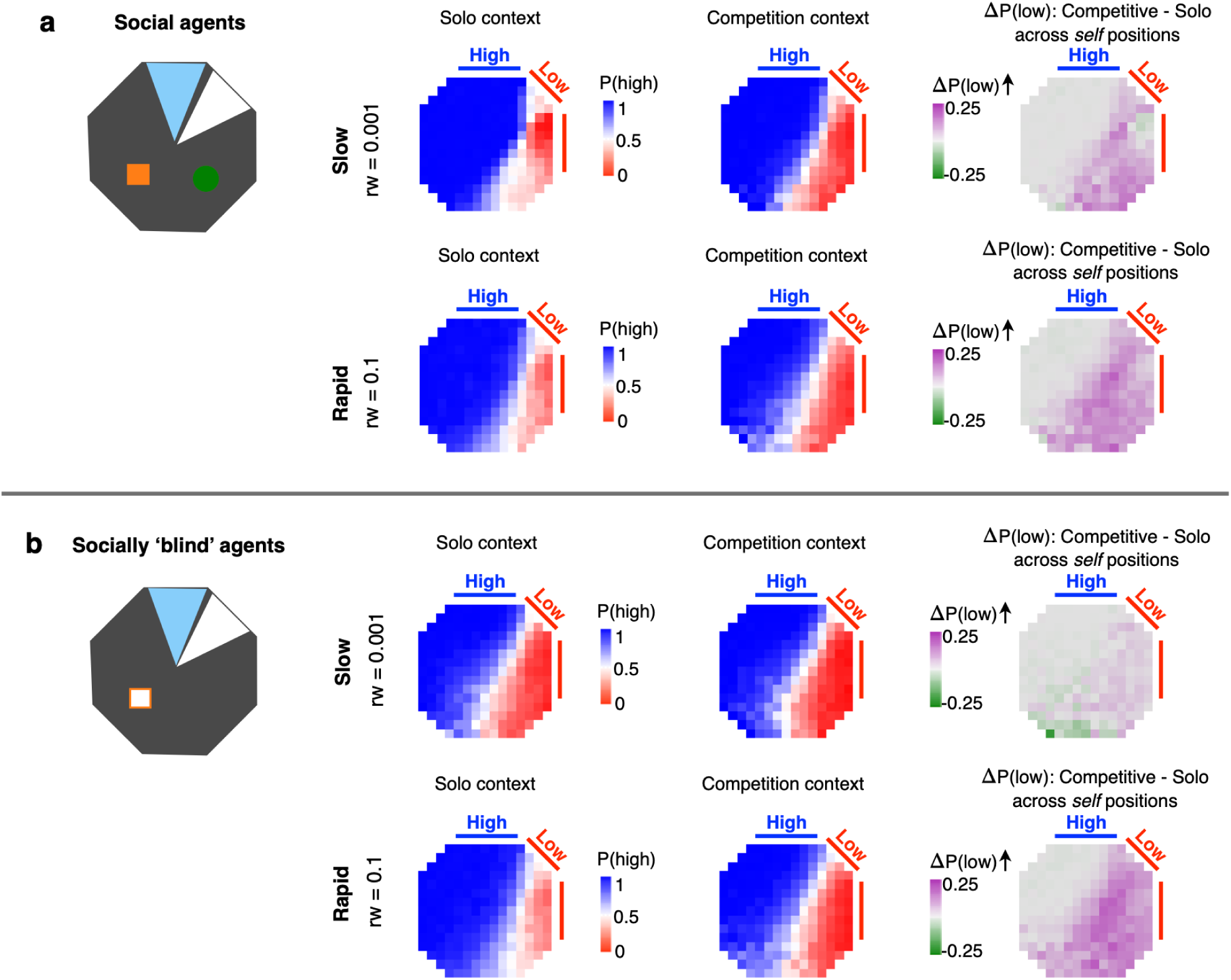
Self-position-specific preference shifts in competing DQN agents. **a**, For ‘social’ DQN agents, preference shifts across contexts varied as a function of self positions in the arena, for both slow learning (top row, rw = 0.001) and rapid learning (bottom row, rw = 0.1). These patterns were similar to mice behaviours (Fig. 2e**,f**). **b**, Similar to **a**, for ‘socially-blind’ agents. Note that with rapid learning, even ‘socially-blind’ agents could display self-position-specific preference shifts (bottom row).

## Statistical Appendix

### Mice Solo Models

For the GLMM exploring the factors influencing Solo decisions, the column definitions were as follows:

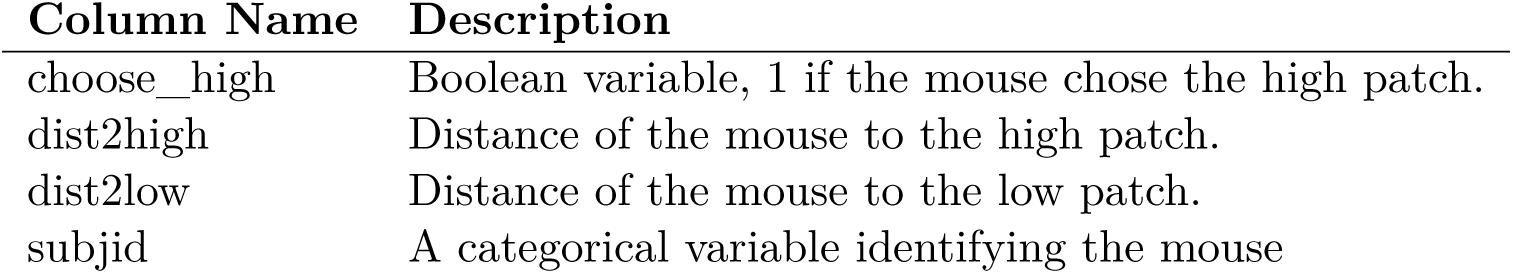

| Column Name | Description |
| --- | --- |
| choose_high | Boolean variable, 1 if the mouse chose the high patch. |
| dist2high | Distance of the mouse to the high patch. |
| dist2low | Distance of the mouse to the low patch. |
| subjid | A categorical variable identifying the mouse |

**Tab. S1:** Mice Solo Model 1.

| AIC | BIC | LogLikelihood | Deviance |
| --- | --- | --- | --- |
| 1.6321e+05 | 1.6329e+05 | -81595 | 1.6319e+05 |

| Name | Estimate | SE | tStat | DF | pValue |
| --- | --- | --- | --- | --- | --- |
| {'(Intercept)'} } | -0.36269 | 0.13422 | -2.7022 | 28979 | 0.0068922 |
| {'dist2high' } } | -1.2268 | 0.13778 | -8.9046 | 28979 | 5.6625e-19 |
| {'dist2low' } } | 5.1992 | 0.16422 | 31.66 | 28979 | 2.6614e-216 |
| {'dist2high:dist2low'} | -1.4618 | 0.11013 | -13.274 | 28979 | 4.3055e-40 |

**Tab. S2:** Mice Solo Model 2.

| AIC | BIC | LogLikelihood | Deviance |
| --- | --- | --- | --- |
| 1.5669e+05 | 1.5677e+05 | -78337 | 1.5667e+05 |

| Name | Estimate | SE | tStat | DF | pValue |
| --- | --- | --- | --- | --- | --- |
| {'(Intercept)'} } | 1.0937 | 0.064651 | 16.918 | 28980 | 6.7781e-64 |
| {'dist2high' } } | -2.5334 | 0.11439 | -22.148 | 28980 | 8.653e-108 |
| {'dist2low' } } | 3.4084 | 0.096151 | 35.448 | 28980 | 1.7588e-269 |

**Tab. S3:** Mice Solo Model 3.

| AIC | BIC | LogLikelihood | Deviance |
| --- | --- | --- | --- |
| 1.4924e+05 | 1.4928e+05 | -74615 | 1.4923e+05 |

| Name | Estimate | SE | tStat | DF | pValue |
| --- | --- | --- | --- | --- | --- |
| {'(Intercept)'} } | -0.52732 | 0.093084 | -5.665 | 28981 | 1.4843e-08 |
| {'dist2low' } } | 2.2228 | 0.076099 | 29.209 | 28981 | 7.1486e-185 |

**Tab. S4:**
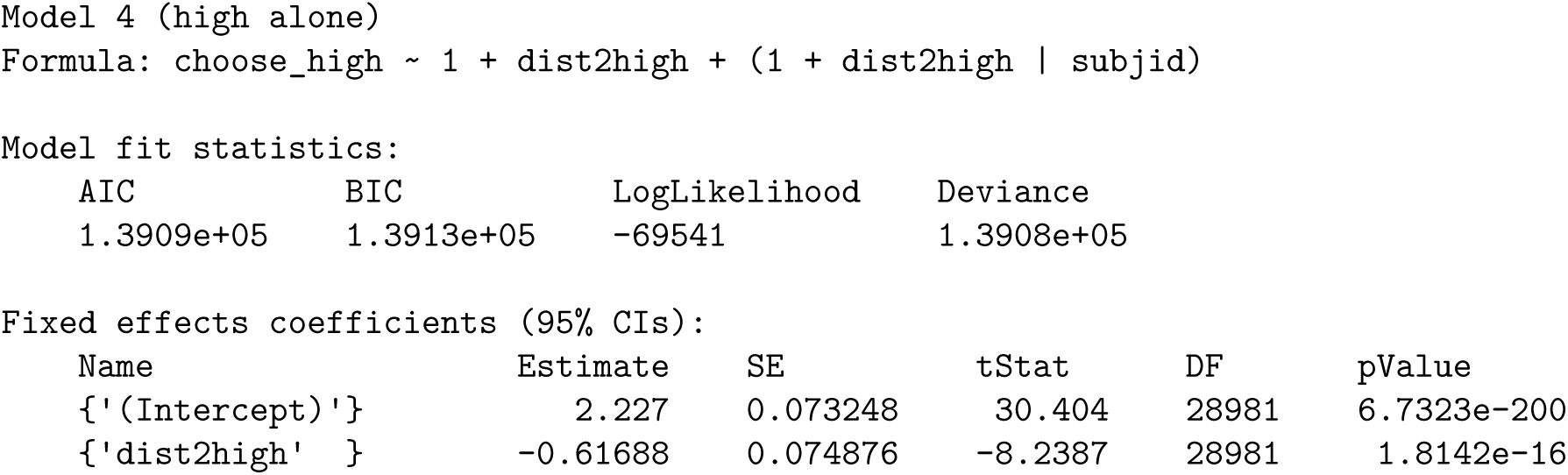
Mice Solo Model 4.

**Tab. S5:**
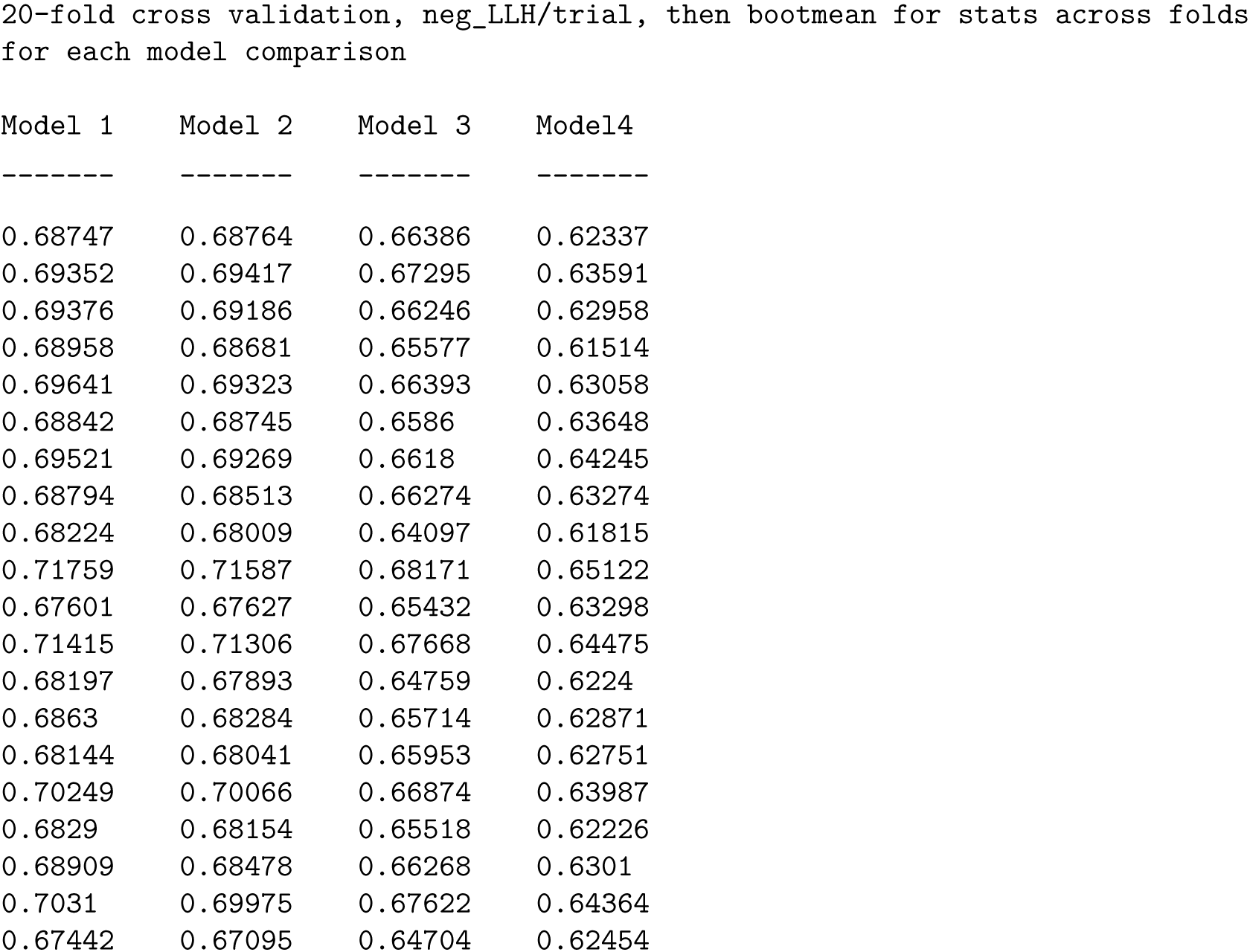
Mice Solo 20-fold cross validation.

| Column Name | Description |
| --- | --- |
| choose_high | Boolean variable 1 if the 'self' mouse chose the high patch. |
| dist2high | Distance of the 'self' mouse to the high patch. |
| dist2low | Distance of the 'self' mouse to the low patch. |
| Social | Boolean variable. 1 if the session was a social session. |
| subjid | A categorical variable identifying the mouse |

Tho column definitions for the GLMM exploring the overall effect of solo vs. social context. The notation for the GLMM formula follows Wilkinson notation as used by the fitglme function in MATLAB.

### Mice Solo vs. Social (without opponent position)

**Tab. S6:**
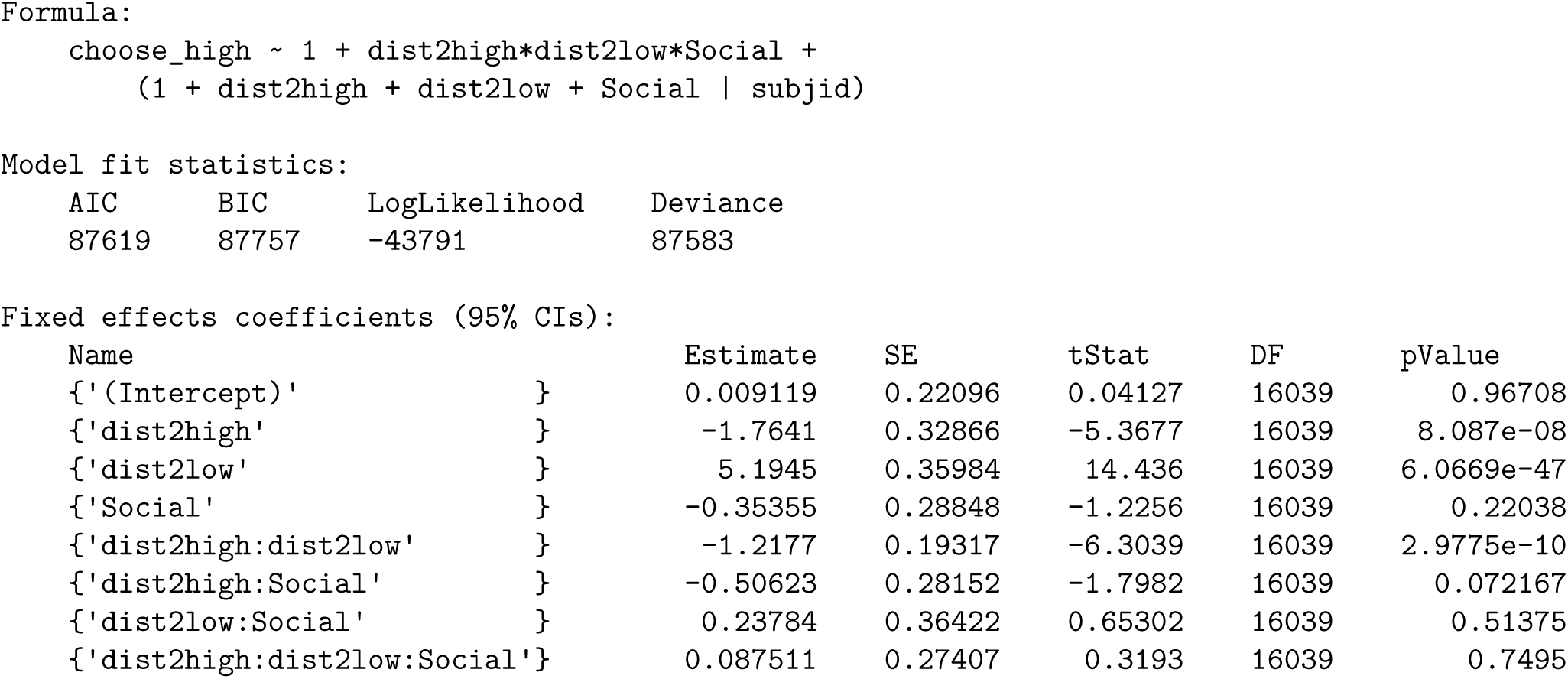
Solo vs. Social: Model 1.

**Tab. S7:**
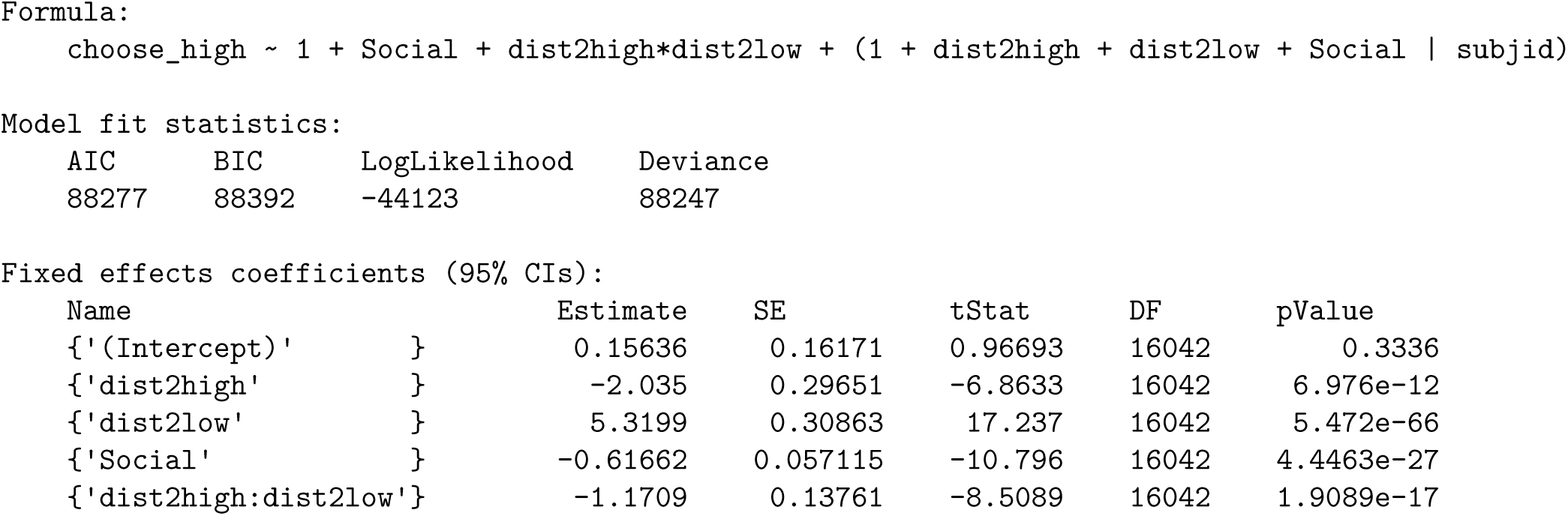
Solo vs. Social: Model 2.

**Tab. S8:**
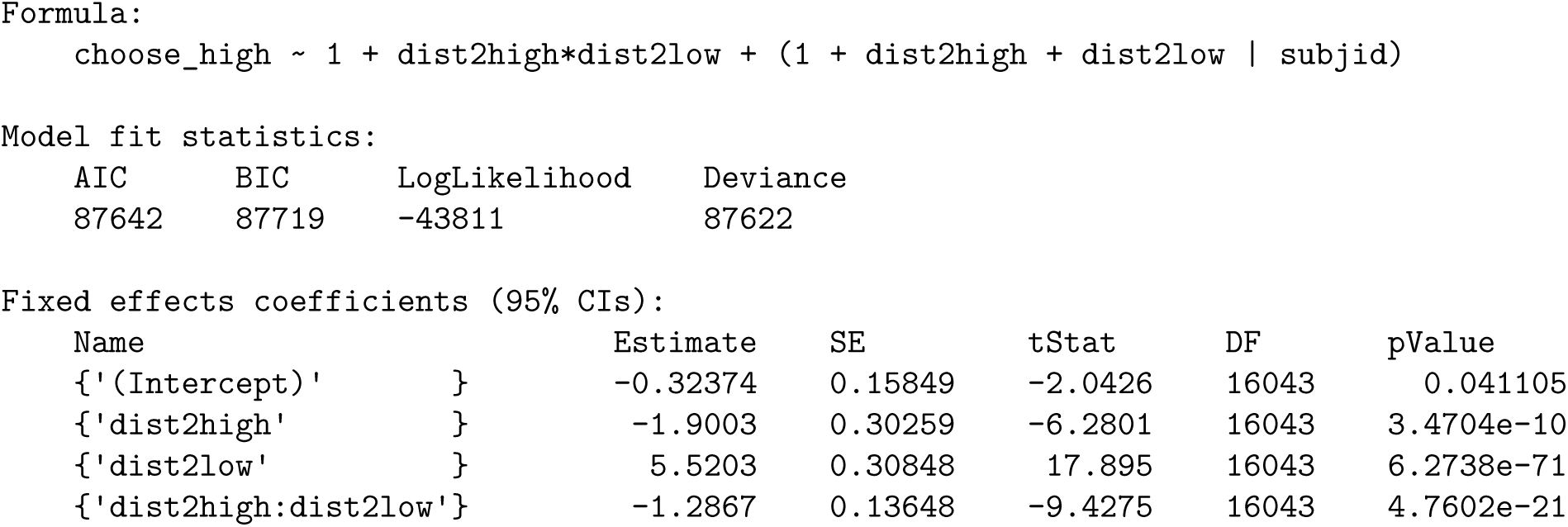
Solo vs. Social: Model 3.

**Tab. S9:**
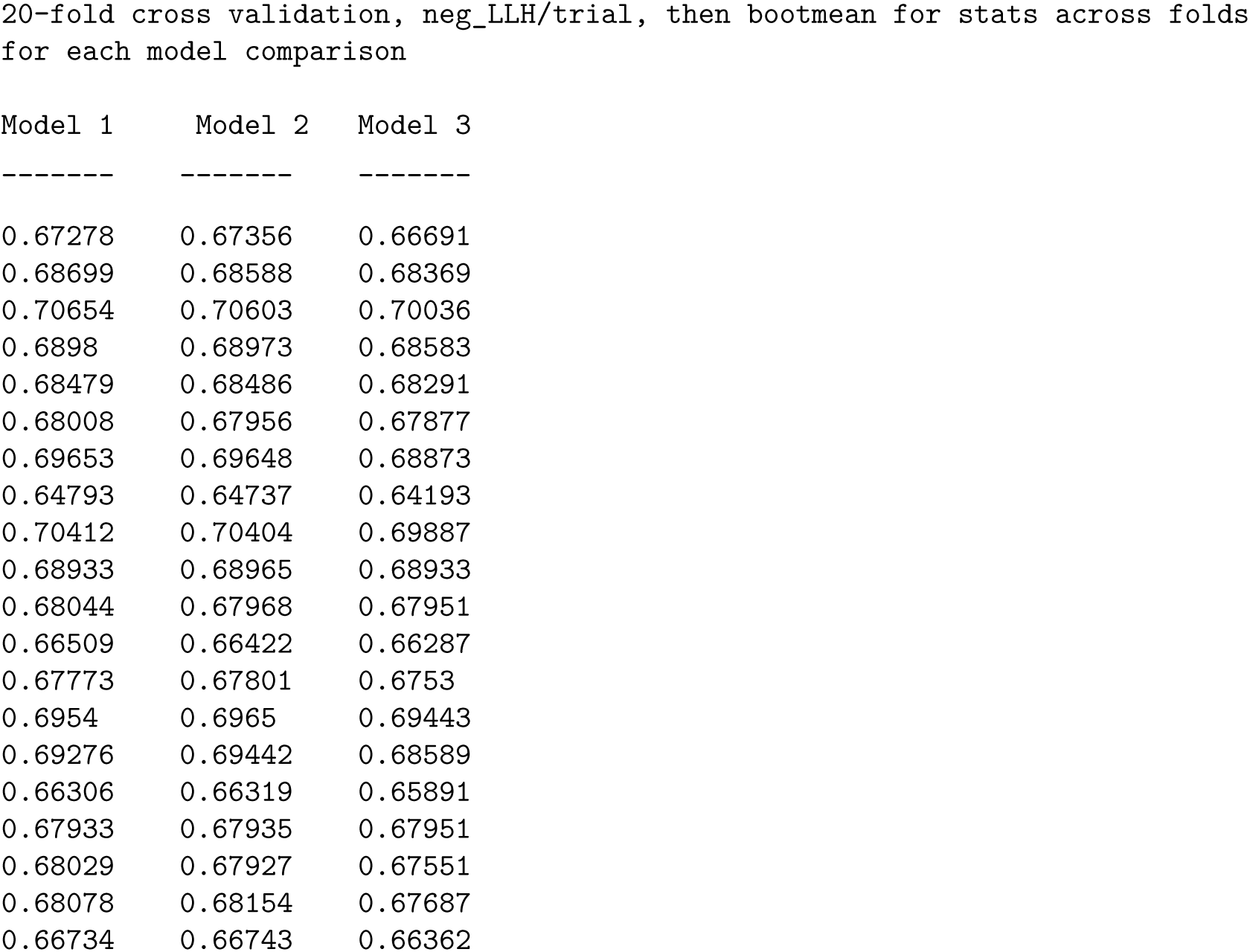
Solo vs. Social: 20-fold cross validation.

### Opponent Effects GLMM

For the GLMM testing the effects of opponent position, the column definitions were as follows:

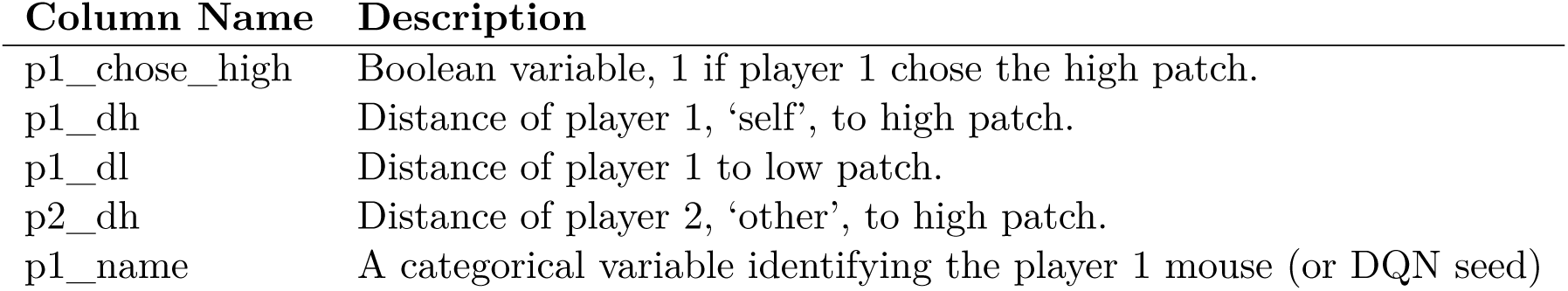

| Column Name | Description |
| --- | --- |
| p1_chose_high | Boolean variable, 1 if player 1 chose the high patch. |
| p1_dh | Distance of player 1, 'self', to high patch. |
| p1_dl | Distance of player 1 to low patch. |
| p2_dh | Distance of player 2, 'other', to high patch. |
| p1_name | A categorical variable identifying the player 1 mouse (or DQN seed) |

The notation for the formula follows Wilkinson notation as used by the MixedModels.jl package in Julia.

### Mice

**Tab. S10:**
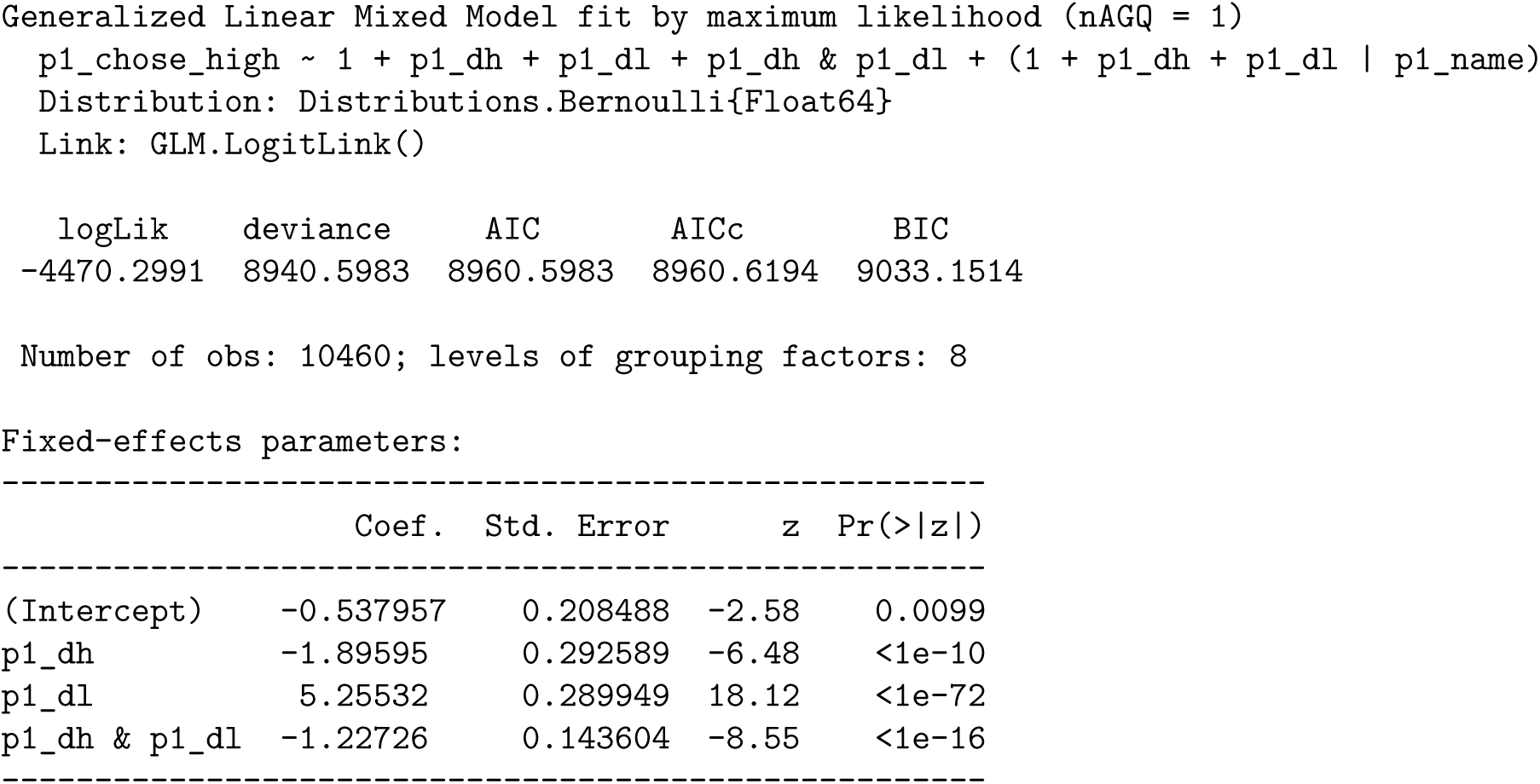
Mice: Model 1.

**Tab. S11:**
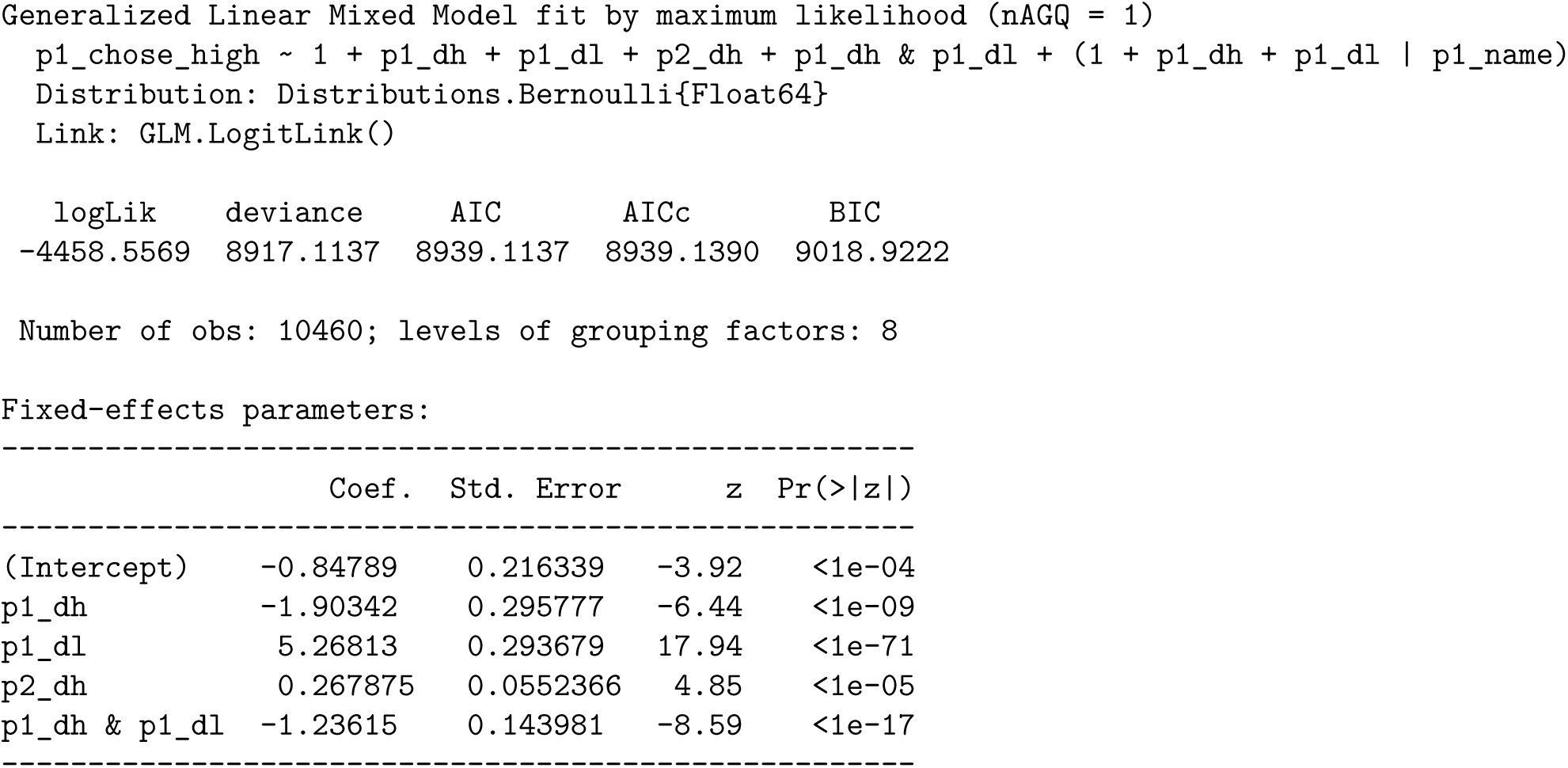
Mice: Model 2.

**Tab. S12:**
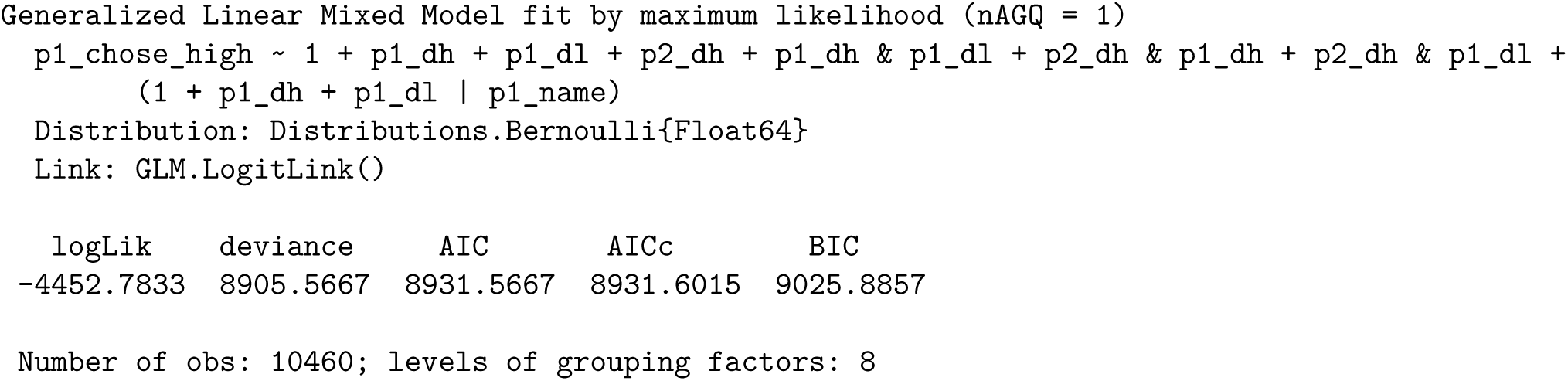
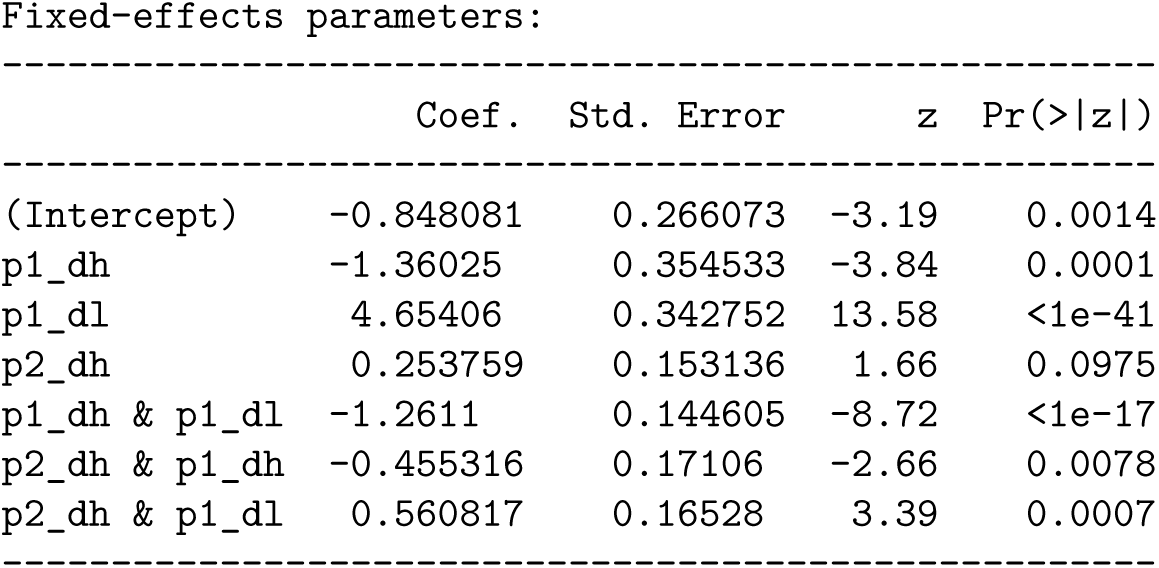
Mice: Model 3.

**Tab. S13:**
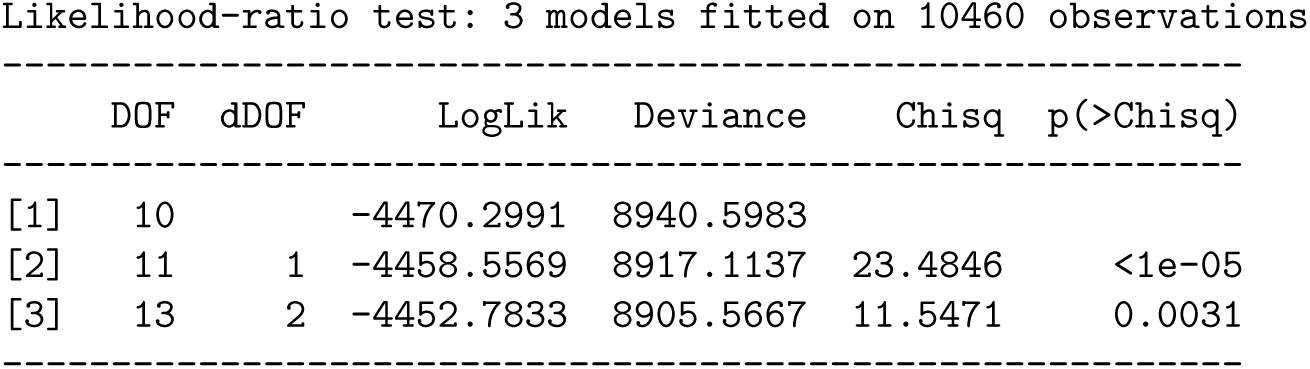
Mice, likelihood ratio tests.

### Social RL

**Tab. S14:**
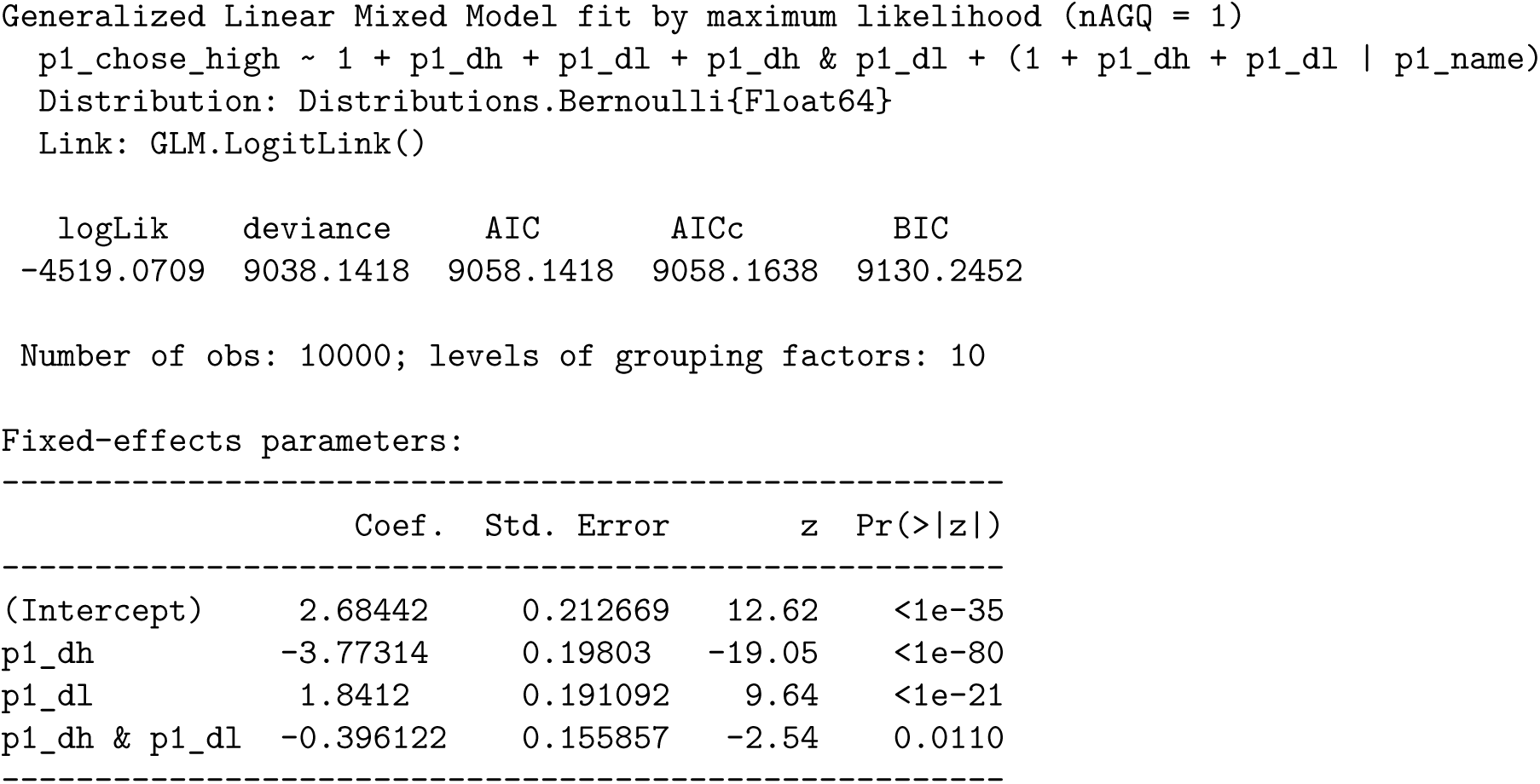
Social RL Model 1.

**Tab. S15:**
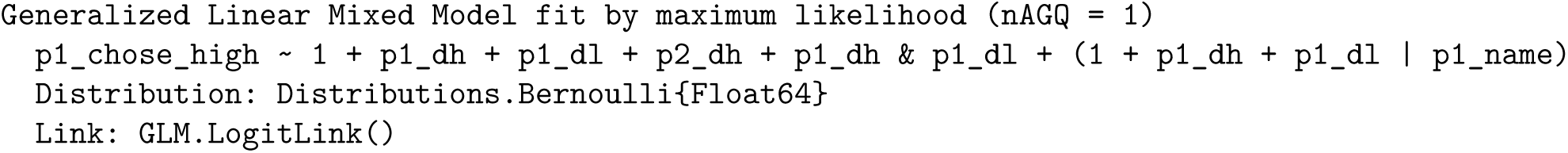
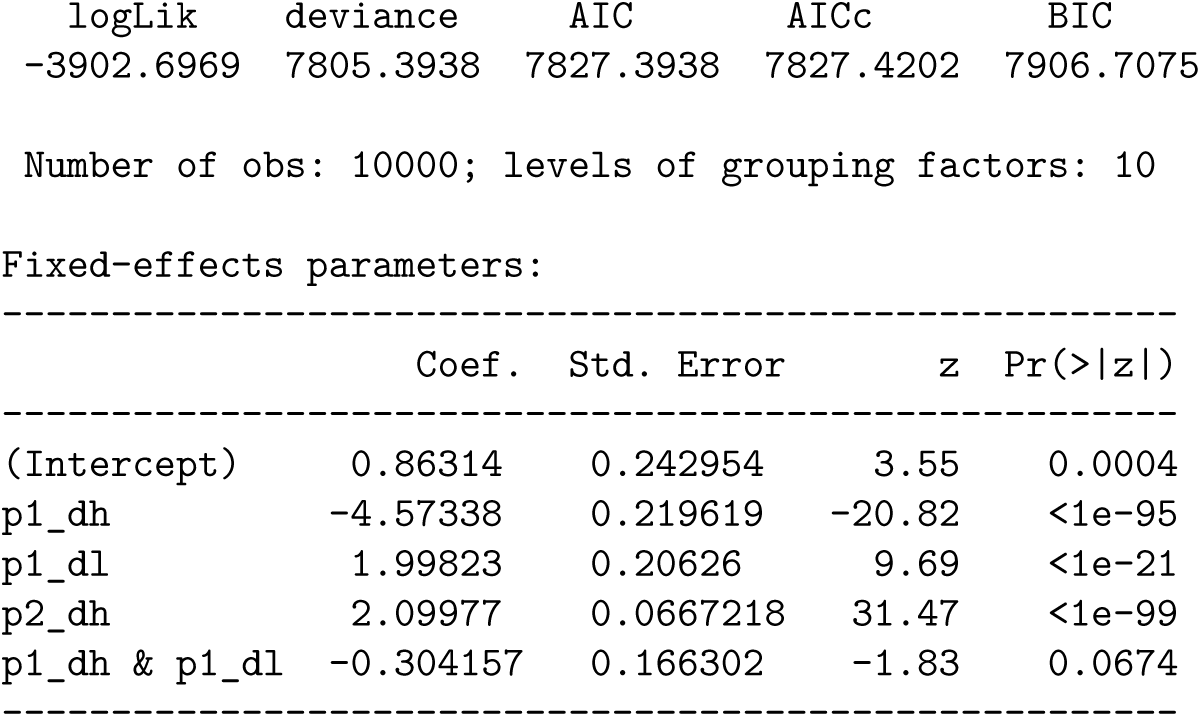
Social RL Model 2.

**Tab. S16:**
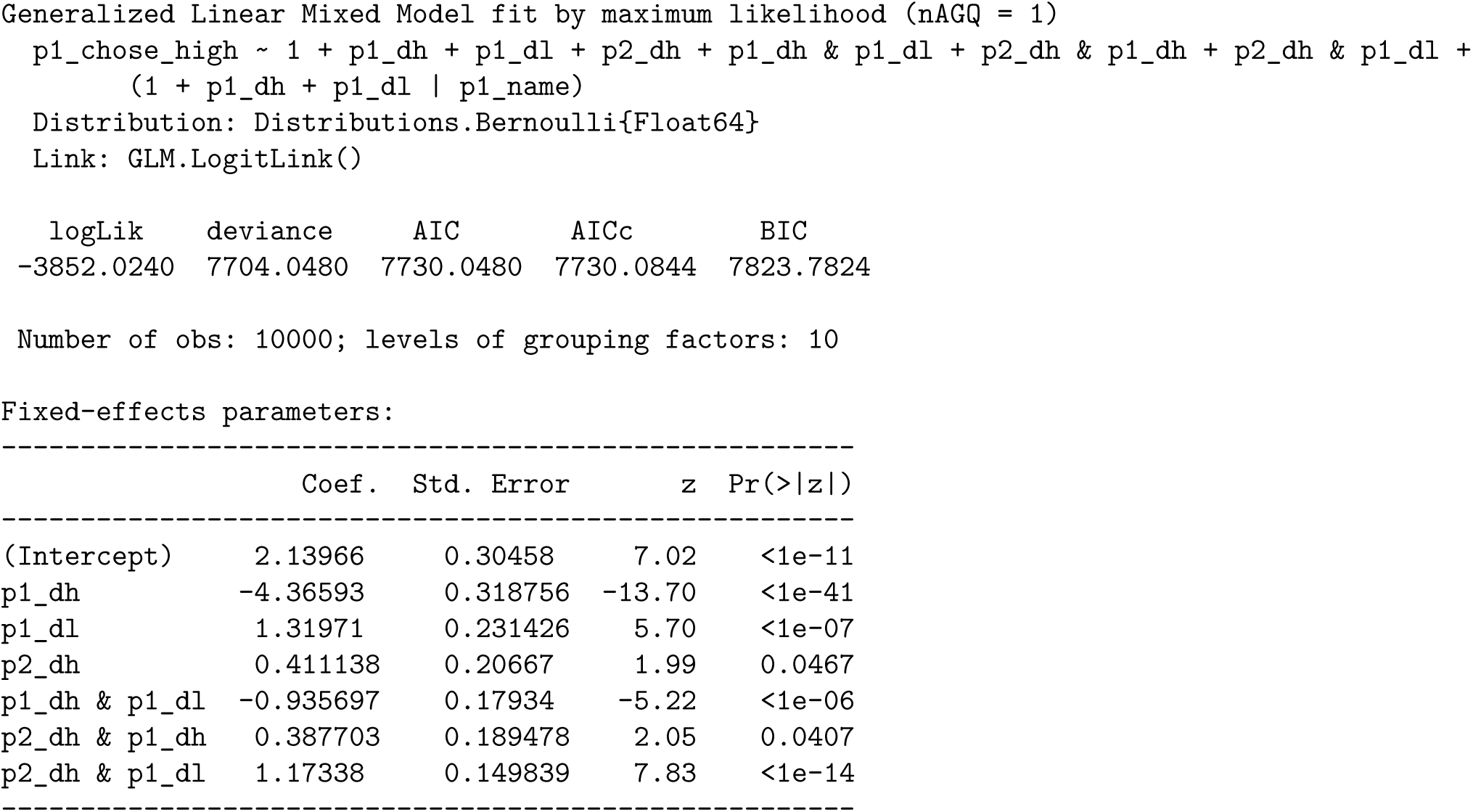
Social RL Model 3.

**Tab. S17:**
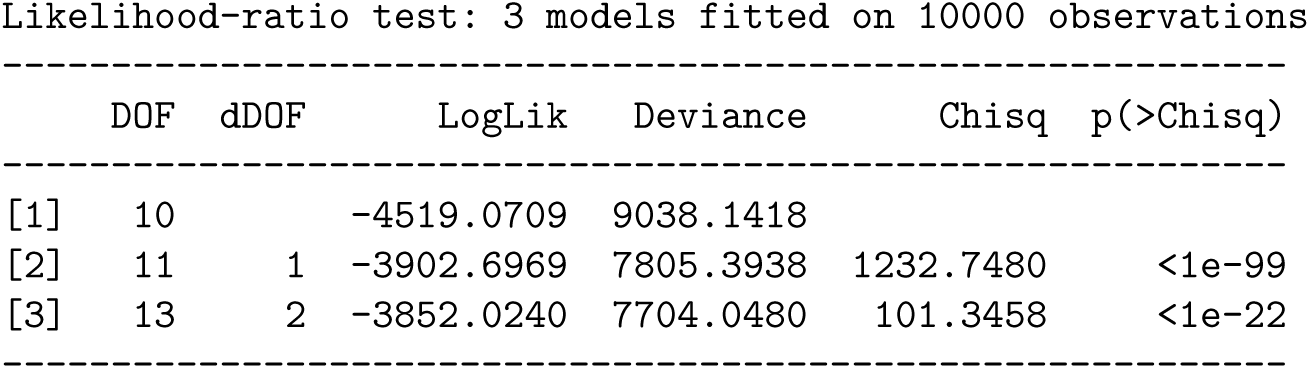
Social RL, likelihood ratio tests.

### Socially-Blind RL

**Tab. S18:**
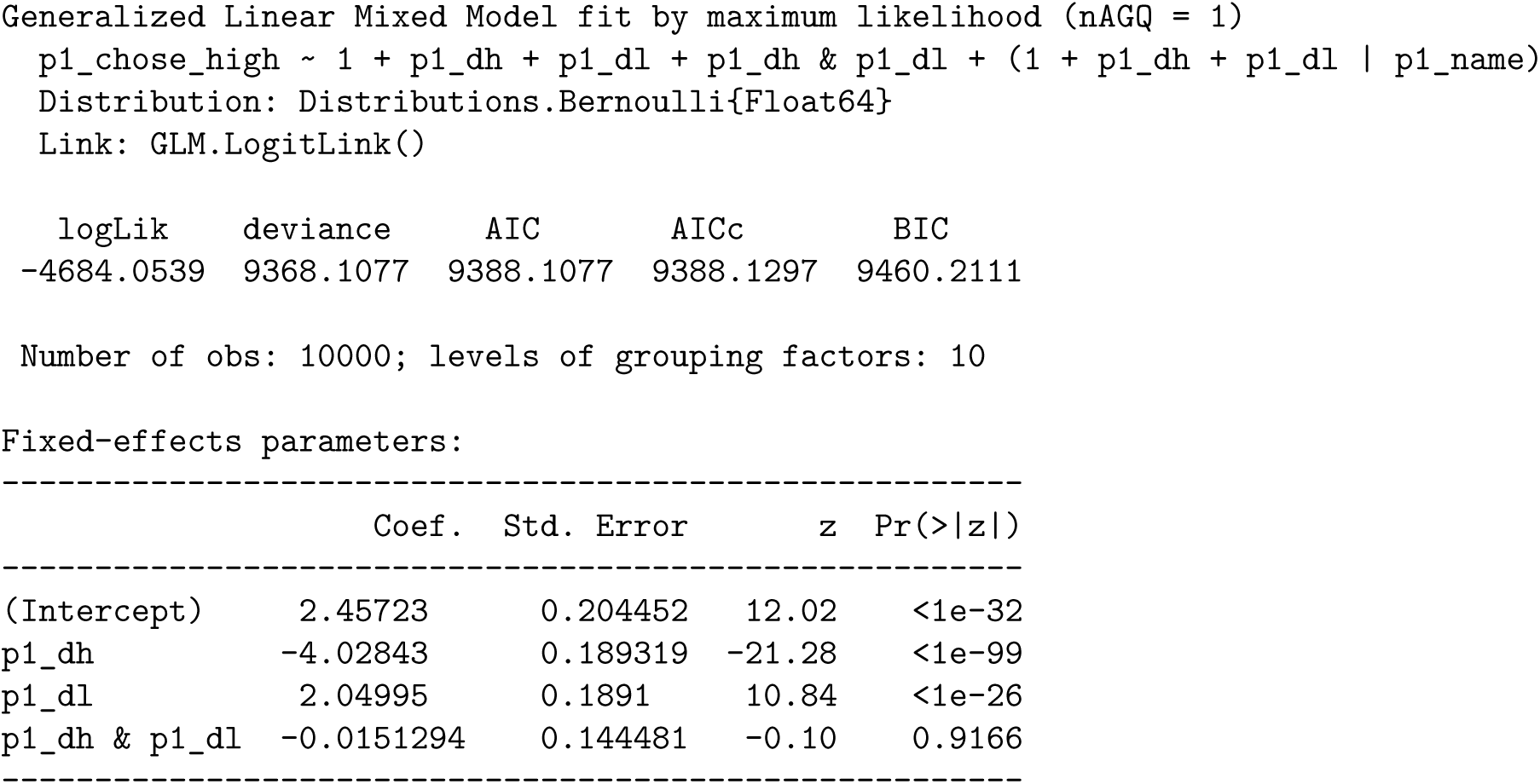
Socially-blind RL Model 1.

**Tab. S19:**
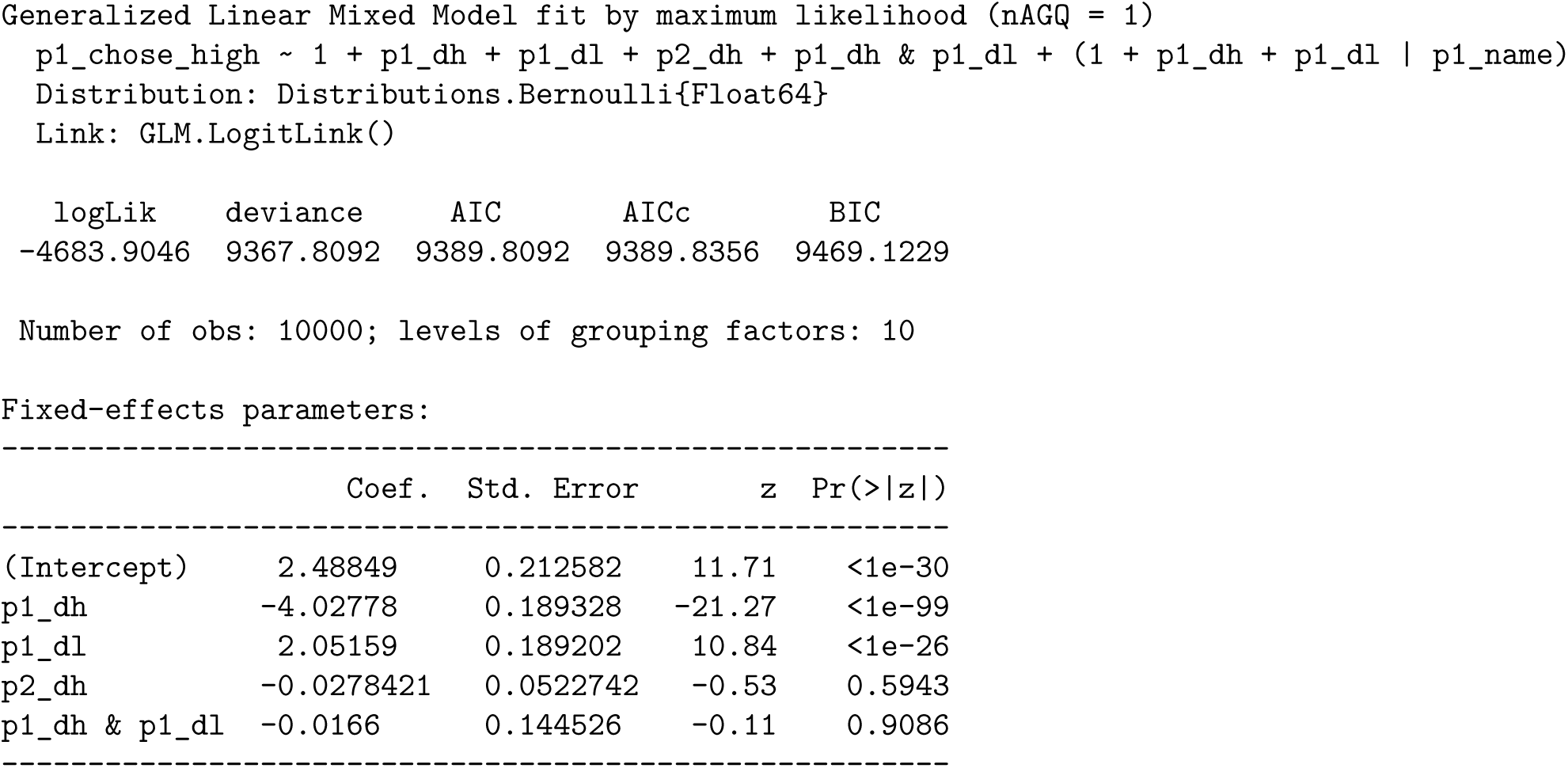
Socially-blind RL Model 2.

**Tab. S20:**
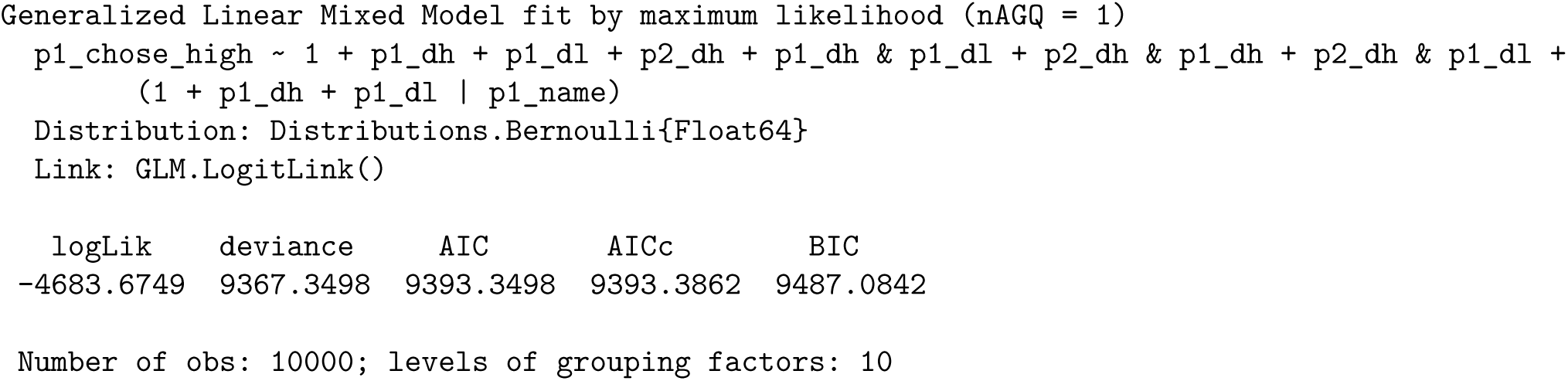
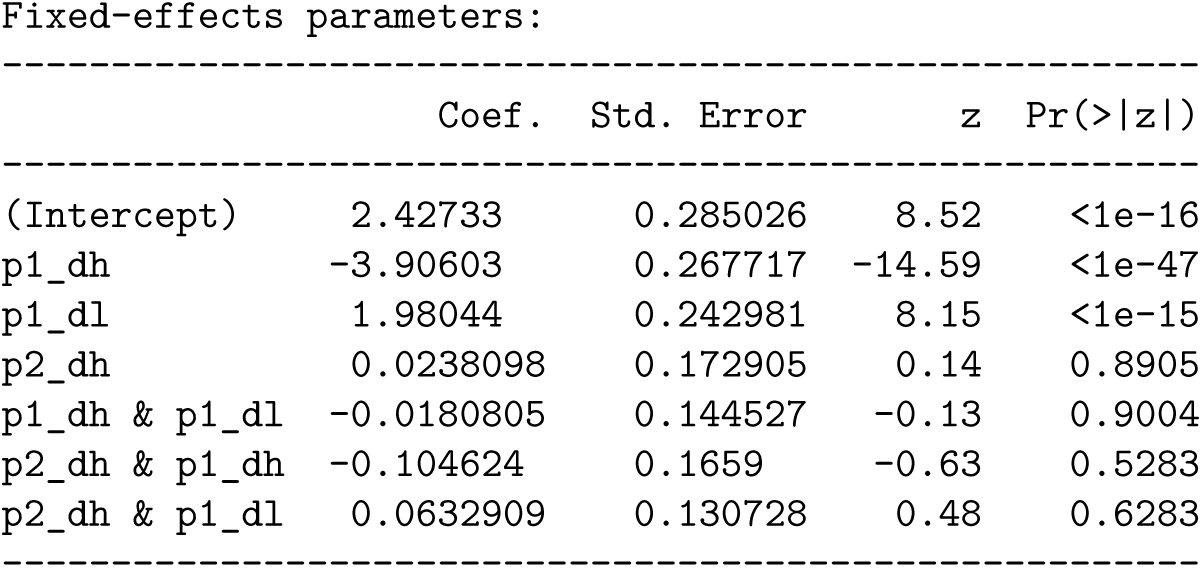
Socially-blind RL Model 3.

**Tab. S21:**
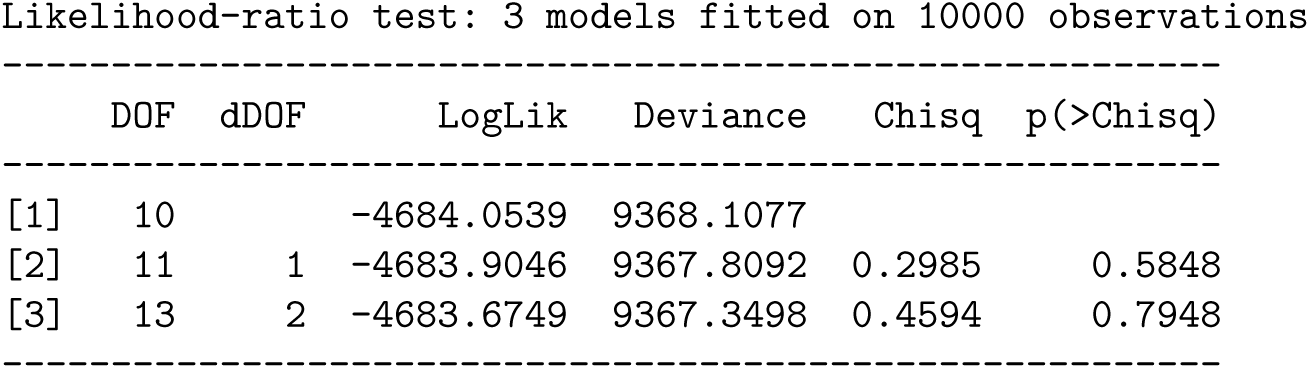
Socially-blind RL, likelihood ratio tests.

### Socially-Blind RL, conditioned on winning

**Tab. S22:**
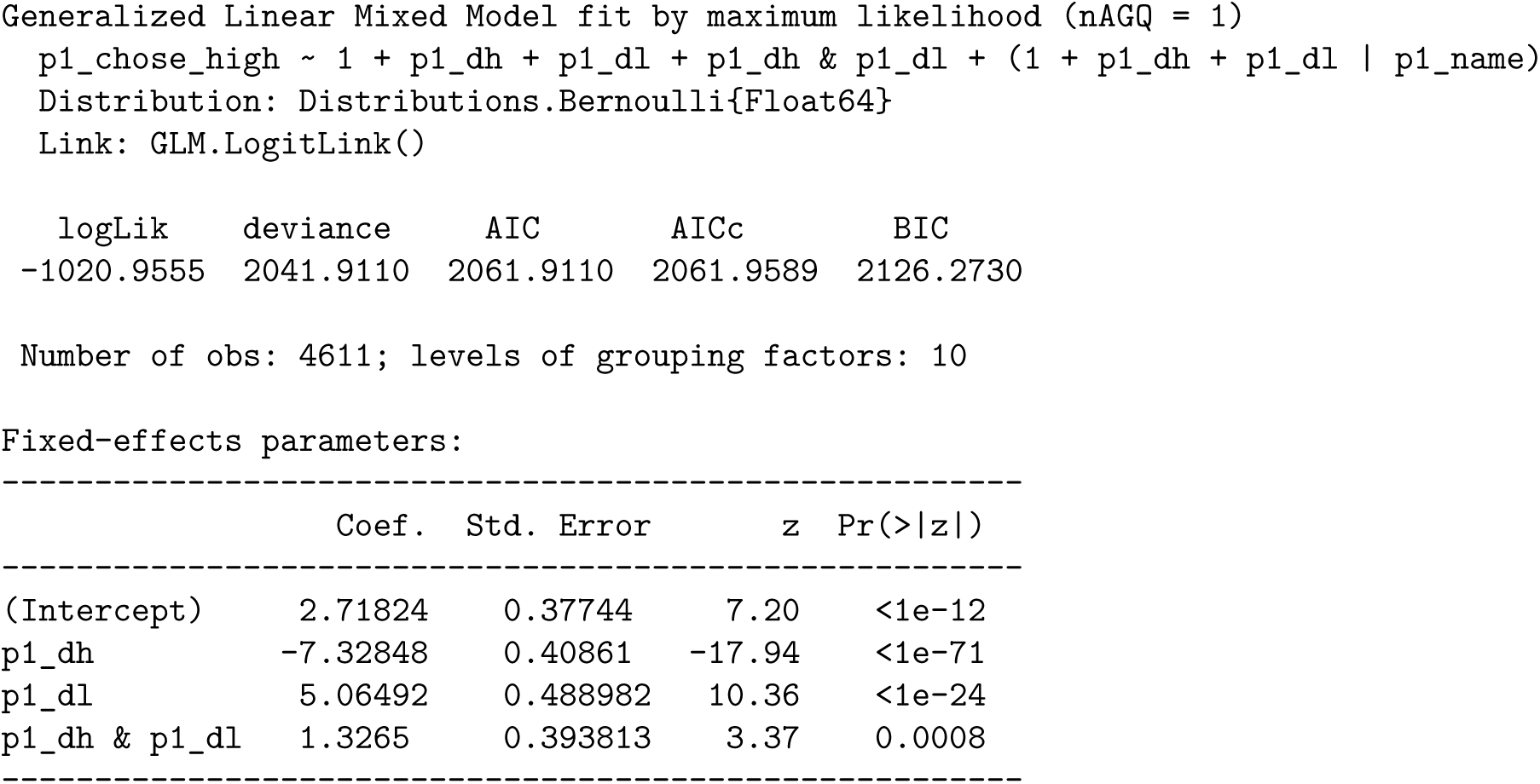
Socially-blind RL, conditioned on winning 1.

**Tab. S23:**
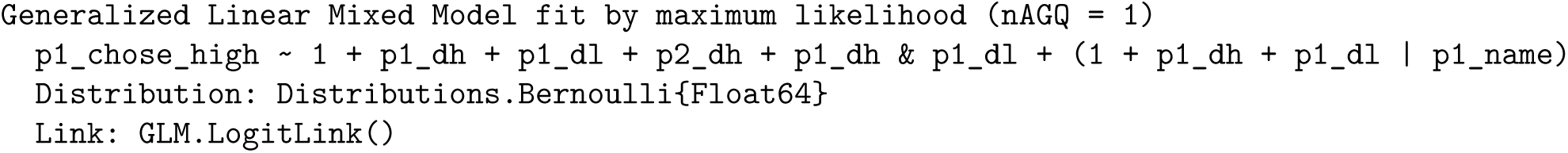
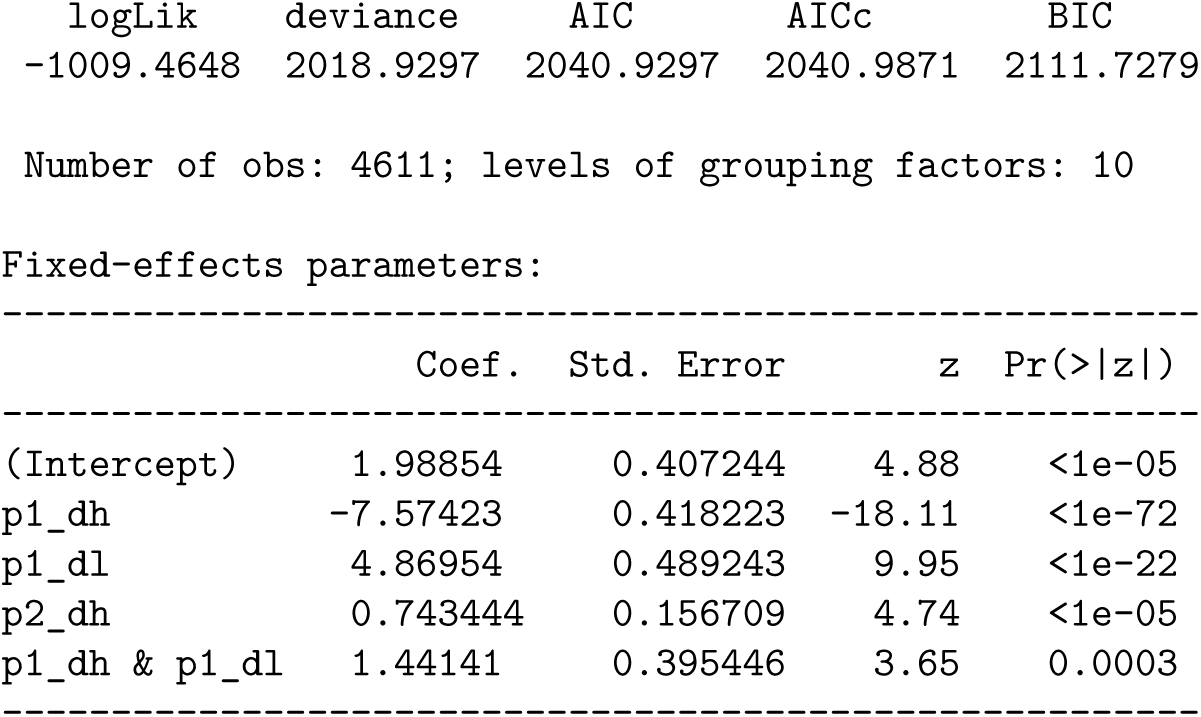
Socially-blind RL, conditioned on winning 2.

**Tab. S24:**
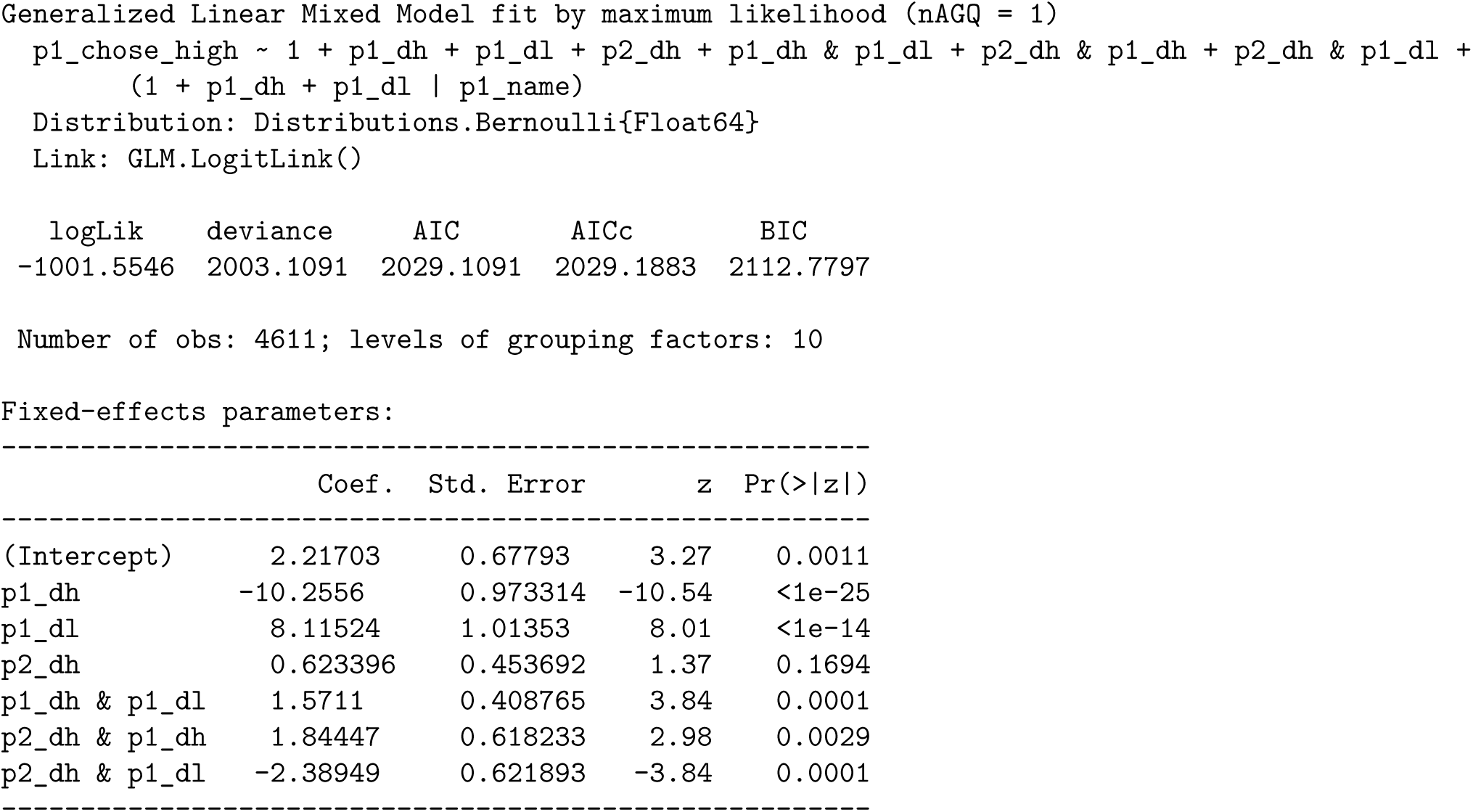
Socially-blind RL, conditioned on winning 3.

**Tab. S25:**
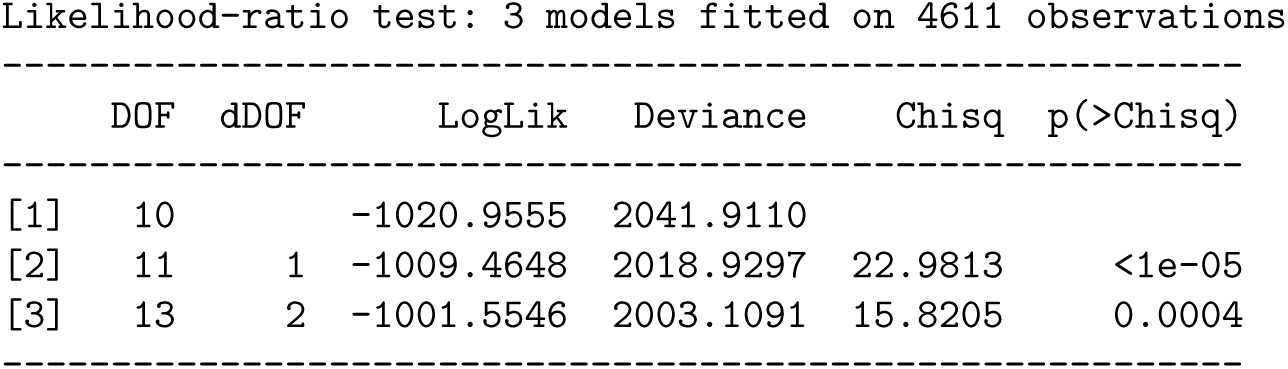
Socially-blind, conditioned on winning: Likelihood Ratio Tests.

## Notes

The authors have no conflict of interest nor financial interests.

https://gitlab.com/sainsbury-wellcome-centre/delab/publications/mice-octagon-2025

### Competing Interest Statement

The authors have declared no competing interest.

### Summary of Updates

Improved flow of text and clarity of figures.

## References

[1] Paul W. Glimcher. The neurobiology of visual-saccadic decision making. Annual Review of Neuroscience, 26:133–179, 2003. doi: 10.1146/annurev.neuro.26.010302.081134.

[2] Karel Svoboda and Nuo Li. Neural mechanisms of movement planning: Motor cortex and beyond. Current Opinion in Neurobiology, 49:33–41, April 2018. doi: 10.1016/j.conb.2017.10.023.

[3] Joshua I. Gold and Michael N. Shadlen. The Neural Basis of Decision Making. Annual Review of Neuroscience, 30(1):535–574, 2007. doi: 10.1146/annurev.neuro.29.051605.113038.

[4] Timothy D. Hanks and Christopher Summerfield. Perceptual Decision Making in Rodents, Monkeys, and Humans. Neuron, 93(1):15–31, January 2017. doi: 10.1016/j.neuron.2016.12.003.

[5] Camillo Padoa-Schioppa. Neurobiology of Economic Choice: A Good-Based Model. Annual Review of Neuroscience, 34(1):333–359, July 2011. doi: 10.1146/annurev-neuro-061010-113648.

[6] Roger Ratcliff. A theory of memory retrieval. Psychological Review, 85(2):59–108, 1978. doi: 10.1037/0033-295X.85.2.59.

[7] Ivana Orsolic, Maxime Rio, Thomas D. Mrsic-Flogel, and Petr Znamenskiy. Mesoscale cortical dynamics reflect the interaction of sensory evidence and temporal expectation during perceptual decision-making. Neuron, 109(11):1861–1875.e10, June 2021. doi: 10.1016/j.neuron.2021.03.031.

[8] Timothy D Hanks, Charles D Kopec, Bingni W Brunton, Chunyu A Duan, Jeffrey C Erlich, and Carlos D Brody. Distinct relationships of parietal and prefrontal cortices to evidence accumulation. Nature, 520(7546):220–223, April 2015. doi: 10.1038/nature14066.

[9] Andrei Khilkevich, Michael Lohse, Ryan Low, Ivana Orsolic, Tadej Bozic, Paige Windmill, and Thomas D. Mrsic-Flogel. Brain-wide dynamics linking sensation to action during decisionmaking. Nature, 634(8035):890–900, October 2024. doi: 10.1038/s41586-024-07908-w.

[10] Daniel Kahneman and Amos Tversky. Prospect Theory: An Analysis of Decision under Risk. Econometrica, 47(2):263, March 1979. doi: 10.2307/1914185.

[11] Michael L. Platt and Paul W. Glimcher. Neural correlates of decision variables in parietal cortex. Nature, 400(6741):233–238, July 1999. doi: 10.1038/22268.

[12] Camillo Padoa-Schioppa and John A. Assad. Neurons in the orbitofrontal cortex encode economic value. Nature, 441(7090):223–226, May 2006. doi: 10.1038/nature04676.

[13] Xinying Cai and Camillo Padoa-Schioppa. Neuronal evidence for good-based economic decisions under variable action costs. Nature Communications, 10(1):393, January 2019. doi: 10.1038/s41467-018-08209-3.

[14] Matthew Rabin and Richard H Thaler. Anomalies: Risk Aversion. Journal of Economic Perspectives, 15(1):219–232, February 2001. doi: 10.1257/jep.15.1.219.

[15] Luc-Alain Giraldeau and Thomas Caraco. Social Foraging Theory. Monographs in Behavior and Ecology. Princeton University Press, Princeton, New Jersey, 2000.

[16] Dougal G. R. Tervo, Mikhail Proskurin, Maxim Manakov, Mayank Kabra, Alison Vollmer, Kristin Branson, and Alla Y. Karpova. Behavioral Variability through Stochastic Choice and Its Gating by Anterior Cingulate Cortex. Cell, 159(1):21–32, September 2014. doi: 10.1016/j.cell.2014.08.037.

[17] Keren Haroush and Ziv M. Williams. Neuronal Prediction of Opponent’s Behavior during Cooperative Social Interchange in Primates. Cell, 160(6):1233–1245, March 2015. doi: 10.1016/j.cell.2015.01.045.

[18] Wei Song Ong, Seth Madlon-Kay, and Michael L. Platt. Neuronal correlates of strategic cooperation in monkeys. Nature Neuroscience, 24(1):116–128, January 2021. doi: 10.1038/s41593-020-00746-9.

[19] Dominic J Barraclough, Michelle L Conroy, and Daeyeol Lee. Prefrontal cortex and decision making in a mixed-strategy game. Nature Neuroscience, 7(4):404–410, April 2004. doi: 10.1038/nn1209.

[20] Michael C. Dorris and Paul W. Glimcher. Activity in Posterior Parietal Cortex Is Correlated with the Relative Subjective Desirability of Action. Neuron, 44(2):365–378, October 2004. doi: 10.1016/j.neuron.2004.09.009.

[21] John Von Neumann and Oskar Morgenstern. Theory of Games and Economic Behavior. Princeton University press, Princeton (N.J.), 3rd ed edition, 1953.

[22] Colin Camerer. Behavioral Game Theory: Experiments in Strategic Interaction. The Roundtable Series in Behavioral Economics. Russell Sage Foundation ; Princeton University Press, New York, N.Y. : Princeton, N.J, 2003.

[23] John F. Nash. Equilibrium points in *n* -person games. Proceedings of the National Academy of Sciences, 36(1):48–49, January 1950. doi: 10.1073/pnas.36.1.48.

[24] Jennifer L. Hoy, Iryna Yavorska, Michael Wehr, and Cristopher M. Niell. Vision Drives Accurate Approach Behavior during Prey Capture in Laboratory Mice. Current Biology, 26 (22):3046–3052, November 2016. doi: 10.1016/j.cub.2016.09.009.

[25] Huai-Ti Lin and Anthony Leonardo. Heuristic Rules Underlying Dragonfly Prey Selection and Interception. Current Biology, 27(8):1124–1137, April 2017. doi: 10.1016/j.cub.2017.03.010.

[26] Philip Coen, Jan Clemens, Andrew J. Weinstein, Diego A. Pacheco, Yi Deng, and Mala Murthy. Dynamic sensory cues shape song structure in Drosophila. Nature, 507(7491):233–237, March 2014. doi: 10.1038/nature13131.

[27] Noga Zilkha, Silvia Gabriela Chuartzman, Yizhak Sofer, Yefim Pen, Meghan Cum, Avi Mayo, Uri Alon, and Tali Kimchi. Sex-dependent control of pheromones on social organization within groups of wild house mice. Current Biology, 33(8):1407–1420.e4, April 2023. doi: 10.1016/j.cub.2023.02.039.

[28] Joshua P Neunuebel, Adam L Taylor, Ben J Arthur, and Se Roian Egnor. Female mice ultrasonically interact with males during courtship displays. eLife, 4:e06203, May 2015. doi: 10.7554/eLife.06203.

[29] Timothy C. Y. Chan, Douglas S. Fearing, Craig Fernandes, and Stephanie Kovalchik. A Markov process approach to untangling intention versus execution in tennis. Journal of Quantitative Analysis in Sports, 18(2):127–145, June 2022. doi: 10.1515/jqas-2021-0077.

[30] Rui Marcelino, Jaime Sampaio, Guy Amichay, Bruno Gonçalves, Iain D. Couzin, and Máté Nagy. Collective movement analysis reveals coordination tactics of team players in football matches. *Chaos*, Solitons & Fractals, 138:109831, September 2020. doi: 10.1016/j.chaos.2020.109831.

[31] Brian M. Sweis, Samantha V. Abram, Brandy J. Schmidt, Kelsey D. Seeland, Angus W. MacDonald, Mark J. Thomas, and A. David Redish. Sensitivity to “sunk costs” in mice, rats, and humans. Science, 361(6398):178–181, July 2018. doi: 10.1126/science.aar8644.

[32] Jake Ormond and John O’Keefe. Hippocampal place cells have goal-oriented vector fields during navigation. Nature, 607(7920):741–746, July 2022. doi: 10.1038/s41586-022-04913-9.

[33] Mohamady El-Gaby, Adam Loyd Harris, James C. R. Whittington, William Dorrell, Arya Bhomick, Mark E. Walton, Thomas Akam, and Timothy E. J. Behrens. A cellular basis for mapping behavioural structure. Nature, 636(8043):671–680, December 2024. doi: 10.1038/s41586-024-08145-x.

[34] Máté Nagy, Attila Horicsányi, Enikő Kubinyi, Iain D. Couzin, Gábor Vásárhelyi, Andrea Flack, and Tamás Vicsek. Synergistic Benefits of Group Search in Rats. Current Biology, 30 (23):4733–4738.e4, December 2020. doi: 10.1016/j.cub.2020.08.079.

[35] Matthew Rosenberg, Tony Zhang, Pietro Perona, and Markus Meister. Mice in a labyrinth show rapid learning, sudden insight, and efficient exploration. eLife, 10:e66175, July 2021. doi: 10.7554/eLife.66175.

[36] Lucas Pinto, Kanaka Rajan, Brian DePasquale, Stephan Y. Thiberge, David W. Tank, and Carlos D. Brody. Task-Dependent Changes in the Large-Scale Dynamics and Necessity of Cortical Regions. Neuron, 104(4):810–824.e9, November 2019. doi: 10.1016/j.neuron.2019.08.025.

[37] Dmitriy Aronov and David W. Tank. Engagement of Neural Circuits Underlying 2D Spatial Navigation in a Rodent Virtual Reality System. Neuron, 84(2):442–456, October 2014. doi: 10.1016/j.neuron.2014.08.042.

[38] P. Simon, R. Dupuis, and J. Costentin. Thigmotaxis as an index of anxiety in mice. Influence of dopaminergic transmissions. Behavioural Brain Research, 61(1):59–64, March 1994. doi: 10.1016/0166-4328(94)90008-6.

[39] Richard P. Heitz. The speed-accuracy tradeoff: History, physiology, methodology, and behavior. Frontiers in Neuroscience, 8, June 2014. doi: 10.3389/fnins.2014.00150.

[40] Volodymyr Mnih, Koray Kavukcuoglu, David Silver, Andrei A. Rusu, Joel Veness, Marc G. Bellemare, Alex Graves, Martin Riedmiller, Andreas K. Fidjeland, Georg Ostrovski, Stig Petersen, Charles Beattie, Amir Sadik, Ioannis Antonoglou, Helen King, Dharshan Kumaran, Daan Wierstra, Shane Legg, and Demis Hassabis. Human-level control through deep reinforcement learning. Nature, 518(7540):529–533, February 2015. doi: 10.1038/nature14236.

[41] C. K. Machens. Flexible Control of Mutual Inhibition: A Neural Model of Two-Interval Discrimination. Science, 307(5712):1121–1124, February 2005. doi: 10.1126/science.1104171.

[42] Charles D Kopec, Jeffrey C Erlich, Bingni W Brunton, Karl Deisseroth, and Carlos D Brody. Cortical and Subcortical Contributions to Short-Term Memory for Orienting Movements. Neuron, 88(2):367–377, October 2015. doi: 10.1016/j.neuron.2015.08.033.

[43] Kong-Fatt Wong and Xiao-Jing Wang. A Recurrent Network Mechanism of Time Integration in Perceptual Decisions. Journal of Neuroscience, 26(4):1314–1328, January 2006. doi: 10.1523/JNEUROSCI.3733-05.2006.

[44] Vivek H. Sridhar, Liang Li, Dan Gorbonos, Máté Nagy, Bianca R. Schell, Timothy Sorochkin, Nir S. Gov, and Iain D. Couzin. The geometry of decision-making in individuals and collectives. Proceedings of the National Academy of Sciences, 118(50):e2102157118, December 2021. doi: 10.1073/pnas.2102157118.

[45] Daniel J. O’Shea, Lea Duncker, Werapong Goo, Xulu Sun, Saurabh Vyas, Eric M. Trautmann, Ilka Diester, Charu Ramakrishnan, Karl Deisseroth, Maneesh Sahani, and Krishna V. Shenoy. Direct neural perturbations reveal a dynamical mechanism for robust computation. bioRxiv, December 2022. doi: 10.1101/2022.12.16.520768.

[46] Philip Shamash, Sarah F. Olesen, Panagiota Iordanidou, Dario Campagner, Nabhojit Banerjee, and Tiago Branco. Mice learn multi-step routes by memorizing subgoal locations. Nature Neuroscience, 24(9):1270–1279, September 2021. doi: 10.1038/s41593-021-00884-8.

[47] Carl D Holmgren, Paul Stahr, Damian J Wallace, Kay-Michael Voit, Emily J Matheson, Juergen Sawinski, Giacomo Bassetto, and Jason Nd Kerr. Visual pursuit behavior in mice maintains the pursued prey on the retinal region with least optic flow. eLife, 10:e70838, October 2021. doi: 10.7554/eLife.70838.

[48] David R. Euston and Bruce L. McNaughton. Apparent Encoding of Sequential Context in Rat Medial Prefrontal Cortex Is Accounted for by Behavioral Variability. Journal of Neuroscience, 26(51):13143–13155, December 2006. doi: 10.1523/JNEUROSCI.3803-06.2006.

[49] Nancy Padilla-Coreano, Kanha Batra, Makenzie Patarino, Zexin Chen, Rachel R. Rock, Ruihan Zhang, Sébastien B. Hausmann, Javier C. Weddington, Reesha Patel, Yu E. Zhang, Hao-Shu Fang, Srishti Mishra, Deryn O. LeDuke, Jasmin Revanna, Hao Li, Matilde Borio, Rachelle Pamintuan, Aneesh Bal, Laurel R. Keyes, Avraham Libster, Romy Wichmann, Fergil Mills, Felix H. Taschbach, Gillian A. Matthews, James P. Curley, Ila R. Fiete, Cewu Lu, and Kay M. Tye. Cortical ensembles orchestrate social competition through hypothalamic outputs. Nature, 603(7902):667–671, March 2022. doi: 10.1038/s41586-022-04507-5.

[50] S. William Li, Omer Zeliger, Leah Strahs, Raymundo Báez-Mendoza, Lance M. Johnson, Aidan McDonald Wojciechowski, and Ziv M. Williams. Frontal neurons driving competitive behaviour and ecology of social groups. Nature, 603(7902):661–666, March 2022. doi: 10.1038/s41586-021-04000-5.

[51] Tingting Zhou, Hong Zhu, Zhengxiao Fan, Fei Wang, Yang Chen, Hexing Liang, Zhongfei Yang, Lu Zhang, Longnian Lin, Yang Zhan, Zheng Wang, and Hailan Hu. History of winning remodels thalamo-PFC circuit to reinforce social dominance. Science, 357(6347):162–168, July 2017. doi: 10.1126/science.aak9726.

[52] Lyle Kingsbury, Shan Huang, Jun Wang, Ken Gu, Peyman Golshani, Ye Emily Wu, and Weizhe Hong. Correlated Neural Activity and Encoding of Behavior across Brains of Socially Interacting Animals. Cell, 178(2):429–446.e16, July 2019. doi: 10.1016/j.cell.2019.05.022.

[53] Oriol Vinyals, Igor Babuschkin, Wojciech M. Czarnecki, Michaël Mathieu, Andrew Dudzik, Junyoung Chung, David H. Choi, Richard Powell, Timo Ewalds, Petko Georgiev, Junhyuk Oh, Dan Horgan, Manuel Kroiss, Ivo Danihelka, Aja Huang, Laurent Sifre, Trevor Cai, John P. Agapiou, Max Jaderberg, Alexander S. Vezhnevets, Rémi Leblond, Tobias Pohlen, Valentin Dalibard, David Budden, Yury Sulsky, James Molloy, Tom L. Paine, Caglar Gulcehre, Ziyu Wang, Tobias Pfaff, Yuhuai Wu, Roman Ring, Dani Yogatama, Dario Wünsch, Katrina McKinney, Oliver Smith, Tom Schaul, Timothy Lillicrap, Koray Kavukcuoglu, Demis Hassabis, Chris Apps, and David Silver. Grandmaster level in StarCraft II using multi-agent reinforcement learning. Nature, 575(7782):350–354, November 2019. doi: 10.1038/s41586-019-1724-z.

[54] Alex T. Piet, Jeffrey C. Erlich, Charles D. Kopec, and Carlos D. Brody. Rat Prefrontal Cortex Inactivations during Decision Making Are Explained by Bistable Attractor Dynamics. Neural Computation, 29(11):2861–2886, August 2017. doi: 10.1162/neco_a_01005.

[55] Chaofei Bao, Xiaoyue Zhu, Joshua Mōller-Mara, Jingjie Li, Sylvain Dubroqua, and Jeffrey C. Erlich. The rat frontal orienting field dynamically encodes value for economic decisions under risk. Nature Neuroscience, 26(11):1942–1952, November 2023. doi: 10.1038/s41593-023-01461-x.

[56] Siddharth Chaturvedi, Ahmed El-Gazzar, and Marcel Van Gerven. A dynamical systems approach to optimal foraging. PLOS Complex Systems, 1(3):e0000018, November 2024. doi: 10.1371/journal.pcsy.0000018.

[57] Mengyu Liu, Aditya Nair, Nestor Coria, Scott W. Linderman, and David J. Anderson. Encoding of female mating dynamics by a hypothalamic line attractor. Nature, 634(8035):901–909, October 2024. doi: 10.1038/s41586-024-07916-w.

[58] Dan Gorbonos, Nir S. Gov, and Iain D. Couzin. Geometrical Structure of Bifurcations during Spatial Decision-Making. PRX Life, 2(1):013008, February 2024. doi: 10.1103/PRXLife.2.013008.

[59] Aditya Nair, Tomomi Karigo, Bin Yang, Surya Ganguli, Mark J. Schnitzer, Scott W. Linderman, David J. Anderson, and Ann Kennedy. An approximate line attractor in the hypothalamus encodes an aggressive state. Cell, 186(1):178–193.e15, January 2023. doi: 10.1016/j.cell.2022.11.027.

[60] Stefan Scherbaum, Simon Frisch, Susanne Leiberg, Steven J. Lade, Thomas Goschke, and Maja Dshemuchadse. Process dynamics in delay discounting decisions: An attractor dynamics approach. Judgment and Decision Making, 11(5):472–495, September 2016. doi: 10.1017/S1930297500004575.

[61] David Laibson. Golden eggs and hyperbolic discounting. The Quarterly Journal of Economics, 112(2):443–478, 1997.

[62] Prabaha Gangopadhyay, Megha Chawla, Olga Dal Monte, and Steve W. C. Chang. Prefrontal– amygdala circuits in social decision-making. Nature Neuroscience, 24(1):5–18, January 2021. doi: 10.1038/s41593-020-00738-9.

[63] Guillaume Hennequin, Tim P. Vogels, and Wulfram Gerstner. Optimal Control of Transient Dynamics in Balanced Networks Supports Generation of Complex Movements. Neuron, 82 (6):1394–1406, June 2014. doi: 10.1016/j.neuron.2014.04.045.

[64] Kerri Smith. Lab mice go wild: Making experiments more natural in order to decode the brain. Nature, 618(7965):448–450, June 2023. doi: 10.1038/d41586-023-01926-w.

[65] Emily Jane Dennis, Ahmed El Hady, Angie Michaiel, Ann Clemens, Dougal R. Gowan Tervo, Jakob Voigts, and Sandeep Robert Datta. Systems Neuroscience of Natural Behaviors in Rodents. The Journal of Neuroscience, 41(5):911–919, February 2021. doi: 10.1523/JNEUROSCI.1877-20.2020.

[66] Talmo D. Pereira, Nathaniel Tabris, Arie Matsliah, David M. Turner, Junyu Li, Shruthi Ravindranath, Eleni S. Papadoyannis, Edna Normand, David S. Deutsch, Z. Yan Wang, Grace C. McKenzie-Smith, Catalin C. Mitelut, Marielisa Diez Castro, John D’Uva, Mikhail Kislin, Dan H. Sanes, Sarah D. Kocher, Samuel S.-H. Wang, Annegret L. Falkner, Joshua W. Shaevitz, and Mala Murthy. SLEAP: A deep learning system for multi-animal pose tracking. Nature Methods, 19(4):486–495, April 2022. doi: 10.1038/s41592-022-01426-1.

[67] Gonçalo Lopes, Niccolò Bonacchi, João Frazão, Joana P. Neto, Bassam V. Atallah, Sofia Soares, Luís Moreira, Sara Matias, Pavel M. Itskov, Patrícia A. Correia, Roberto E. Medina, Lorenza Calcaterra, Elena Dreosti, Joseph J. Paton, and Adam R. Kampff. Bonsai: An event-based framework for processing and controlling data streams. Frontiers in Neuroinformatics, 9, April 2015. doi: 10.3389/fninf.2015.00007.

[68] Alessio Paolo Buccino, Mikkel Elle Lepperød, Svenn-Arne Dragly, Philipp Häfliger, Marianne Fyhn, and Torkel Hafting. Open source modules for tracking animal behavior and closed-loop stimulation based on Open Ephys and Bonsai. Journal of Neural Engineering, 15(5):055002, October 2018. doi: 10.1088/1741-2552/aacf45.

[69] Jessy Lauer, Mu Zhou, Shaokai Ye, William Menegas, Steffen Schneider, Tanmay Nath, Mohammed Mostafizur Rahman, Valentina Di Santo, Daniel Soberanes, Guoping Feng, Venkatesh N. Murthy, George Lauder, Catherine Dulac, Mackenzie Weygandt Mathis, and Alexander Mathis. Multi-animal pose estimation, identification and tracking with DeepLab-Cut. Nature Methods, 19(4):496–504, April 2022. doi: 10.1038/s41592-022-01443-0.

[70] Tristan Walter and Iain D Couzin. TRex, a fast multi-animal tracking system with markerless identification, and 2D estimation of posture and visual fields. eLife, 10:e64000, February 2021. doi: 10.7554/eLife.64000.

[71] Dean Mobbs, Pete C. Trimmer, Daniel T. Blumstein, and Peter Dayan. Foraging for foundations in decision neuroscience: Insights from ethology. Nature Reviews Neuroscience, 19(7): 419–427, July 2018. doi: 10.1038/s41583-018-0010-7.

[72] Patricia L. Lockwood, Matthew A.J. Apps, and Steve W.C. Chang. Is There a ‘Social’ Brain? Implementations and Algorithms. Trends in Cognitive Sciences, 24(10):802–813, October 2020. doi: 10.1016/j.tics.2020.06.011.

[73] Camille Testard, Sébastien Tremblay, Felipe Parodi, Ron W. DiTullio, Arianna Acevedo-Ithier, Kristin L. Gardiner, Konrad Kording, and Michael L. Platt. Neural signatures of natural behaviour in socializing macaques. Nature, 628(8007):381–390, April 2024. doi: 10.1038/s41586-024-07178-6.

[74] Angelo Forli and Michael M. Yartsev. Hippocampal representation during collective spatial behaviour in bats. Nature, 621(7980):796–803, September 2023. doi: 10.1038/s41586-023-06478-7.

[75] Saikat Ray, Itay Yona, Nadav Elami, Shaked Palgi, Kenneth W. Latimer, Bente Jacobsen, Menno P. Witter, Liora Las, and Nachum Ulanovsky. Hippocampal coding of identity, sex, hierarchy, and affiliation in a social group of wild fruit bats. Science, 387(6733):eadk9385, January 2025. doi: 10.1126/science.adk9385.

[76] Lindsay Willmore, Courtney Cameron, John Yang, Ilana B. Witten, and Annegret L. Falkner. Behavioural and dopaminergic signatures of resilience. Nature, 611(7934):124–132, November 2022. doi: 10.1038/s41586-022-05328-2.

[77] Sharon C. Furtak, Christine E. Cho, Kristin M. Kerr, Jennifer L. Barredo, Janelle E. Alleyne, Yolanda R. Patterson, and Rebecca D. Burwell. The Floor Projection Maze: A novel behavioral apparatus for presenting visual stimuli to rats. Journal of Neuroscience Methods, 181 (1):82–88, June 2009. doi: 10.1016/j.jneumeth.2009.04.023.

[78] Sainsbury Wellcome Centre Foraging Behaviour Working Group. Aeon: An open-source platform to study the neural basis of ethological behaviours over naturalistic timescales. Zenodo 10.5281/ZENODO.8411157, October 2023.

[79] Jeff Bezanson, Alan Edelman, Stefan Karpinski, and Viral B. Shah. Julia: A Fresh Approach to Numerical Computing. SIAM Review, 59(1):65–98, January 2017. doi: 10.1137/141000671.

[80] Phillip M. Alday and Douglas Bates. MixedModels.jl. Zenodo 10.5281/ZENODO.596435, February 2025.

[81] Simon Danisch and Julius Krumbiegel. Makie.jl: Flexible high-performance data visualization for Julia. Journal of Open Source Software, 6(65):3349, 2021. doi: 10.21105/joss.03349.

[82] Xavier Glorot and Yoshua Bengio. Understanding the difficulty of training deep feedforward neural networks. Proceedings of the Thirteenth International Conference on Artificial Intelligence and Statistics (AISTATS), 9:249–256, 2010.

